# Repeated Mild Head Injury Establishes a Senescent Cranial Bone Marrow Niche that Impairs Brain Metabolism

**DOI:** 10.64898/2026.01.12.699107

**Authors:** Patrick J. Devlin, Romeesa Khan, Trang H. Do, Bryce E. West, Janelle M. Korf, Gary U. Guzman, Chunfeng Tan, John Ahn, Rene Flores, Micheal E. Maniskas, Sean P. Marrelli, Anna Malovannaya, Erica Underwood, Rodney M. Ritzel

## Abstract

Traumatic brain injury (TBI) of any severity is associated with long-term systemic inflammation and increased risk of peripheral comorbidities, yet the mechanisms driving immune dysregulation and accelerated aging after repeated mild head impacts remain poorly defined. Here, we investigated the acute and chronic effects of repeated mild TBI (rmTBI) on distal and proximal bone marrow compartments in the femur and calvaria, respectively. Using a modified weight-drop mouse model delivering rotational and acceleration–deceleration forces (3 hits/week for up to 16 weeks), rmTBI produced no mortality, skull fracture, hemorrhage, or brain leukocyte infiltration. One day after three consecutive impacts, rmTBI induced robust proliferation of LSK stem/progenitor cells in both femoral and calvarial marrow, evidenced by Ki67 expression, BrdU incorporation, and increased monocyte output. By 8 weeks (24 impacts), injury-induced proliferation subsided and LSK cells exhibited increased senescence-associated β-galactosidase activity and upregulation of tumor suppressor genes. At 16 weeks (48 impacts), LSK populations were depleted at both sites, displaying reduced proliferative capacity, telomere shortening, and pancytopenia in otherwise young adult mice. Calvarial bone marrow cells exposed to rmTBI released a distinct cytokine and proteomic secretome marked by elevated IL-6, suppressed mitochondrial and metabolic signaling, and enhanced DNA repair pathways. Notably, skull-derived secretome factors impaired cortical and hippocampal mitochondrial metabolism, and reduced microglial mitochondrial membrane potential. Together, these findings identify replicative senescence of the brain-adjacent bone marrow niche as an early and progressive consequence of repeated mild head injury, linking rmTBI to long-lasting metabolic dysfunction, impaired immunity, and accelerated aging.

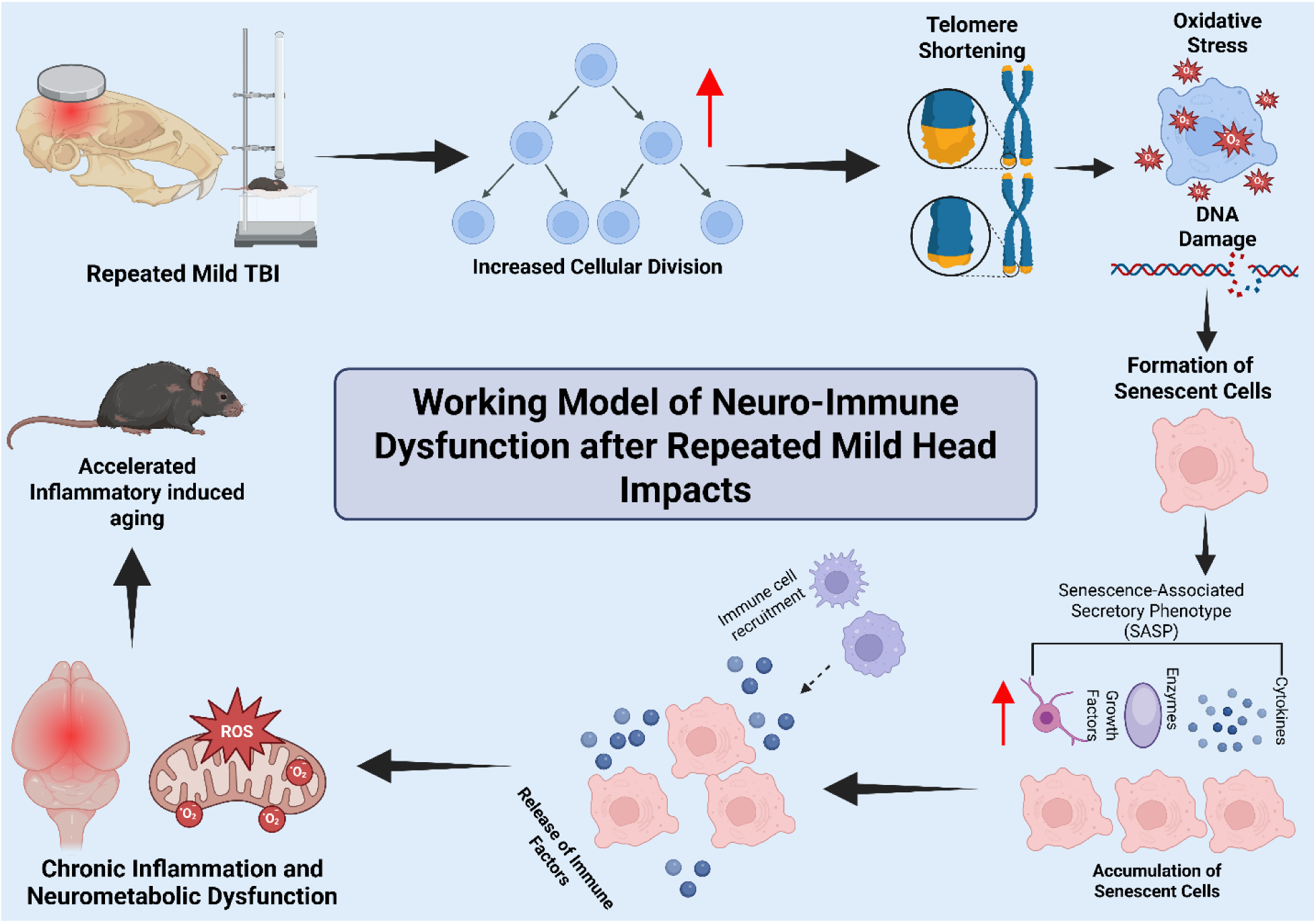

## Introduction

Mild traumatic brain injury (mTBI) accounts for approximately 70 –90% of all reported TBI cases[1] and represents a complex pathophysiological process typically associated with transient neurological, psychological, and cognitive disturbances that may persist long term[2]. mTBI is defined by biomechanical forces applied to the head that induce acute neurological abnormalities, with or without loss of consciousness[3]. Repetitive head impacts are a major contributor to mTBI burden, with sports-related TBIs accounting for ∼6% of all emergency department visits. The repetitive nature of these injuries increases the likelihood of cumulative damage, exacerbating adverse neurological outcomes over time. Indeed, 10–15% of individuals with mTBI develop long-term disabilities or persistent post-concussion symptoms, including cognitive, emotional, and physical impairments[1].

Repeated mild traumatic brain injuries (rmTBI), including sub-concussive impacts commonly experienced in contact sports, are increasingly recognized as drivers of chronic neurological dysfunction. rmTBI is associated with long-term sequelae such as post-concussion syndrome and chronic traumatic encephalopathy (CTE), a progressive neurodegenerative disorder that can emerge years to decades after repeated head trauma[4]. Clinical manifestations may evolve over time, with anxiety and depression appearing earlier, followed by memory deficits, motor dysfunction, and eventual dementia[5]. Both human and animal studies demonstrate that repeated sub-concussive impacts disrupt white matter integrity, reduce cortical thickness, and impair cognitive function[6–9]. Secondary injury mechanisms contributing to these chronic outcomes include neuroinflammation, oxidative stress, and metabolic dysfunction[10]. Despite this growing body of work, the progressive and cumulative effects of rmTBI on systemic immunity- and its contribution to chronic brain inflammation, accelerated aging, and metabolic dysfunction-remain poorly understood.

Neuroinflammation is a central feature of TBI pathology and a key driver of progressive neurodegeneration[11]. Chronic microglial activation is a hallmark of rmTBI, as demonstrated by postmortem neuropathology[12] and translocator protein (TSPO) positron emission tomography (PET) imaging in active collegiate athletes with concussion and former American football players[13]. Microglial dysfunction has been implicated in tau accumulation in human CTE[14]. Experimentally, targeting neuroinflammation through anti-inflammatory strategies or microglial depletion and repopulation improves long-term outcomes following repetitive head impacts[15] and moderate-to-severe TBI[16], underscoring the tight interdependence between neuroinflammation and neurodegeneration.

Less well defined, however, is the role of systemic immune dysregulation in shaping the chronic consequences of rmTBI. Following TBI, activation of the peripheral immune system and sustained release of inflammatory mediators into the circulation may influence the brain through blood–brain barrier (BBB) disruption and leukocyte trafficking[17, 18]. Systemic inflammation can persist for months to years after mTBI and correlates with cognitive impairment and neurological symptoms[19, 20]. Notably, TBI of any severity is associated with an elevated risk of chronic cardiovascular, endocrine, and neurological comorbidities over a 10 - year period, even among younger individuals[21]. These findings highlight the widespread and bidirectional systemic effects of TBI and implicate immune dysfunction as a potential contributor to long-term pathology. Consistent with this concept, we recently demonstrated that a single moderate-to-severe TBI induces long-lasting hematopoietic stem/progenitor cell senescence in the bone marrow, driving chronic systemic inflammation, cognitive dysfunction, and neurodegenerative disease signatures[22]. However, the progressive and cumulative impact of rmTBI on hematopoiesis-particularly within skull-associated bone marrow-remains largely unexplored, despite evidence of chronic leukocyte alterations, shortened leukocyte telomeres, and persistent blood cytokine changes in rmTBI populations[23, 24]. Longitudinal characterization of these processes in humans is hindered by diagnostic ambiguity, injury heterogeneity, comorbidities, attrition, and a lack of reliable biomarkers.

While bone marrow responses to TBI have been primarily studied in long bones such as the femur following moderate-to-severe injury, the calvaria-a flat bone forming the skullcap-has received comparatively little attention. Emerging evidence indicates that calvarial bone marrow cells respond to systemic infection and brain injury by migrating through ossified vascular channels directly into the meninges, bypassing the BBB[25–27]. Importantly, distal femoral and proximal calvarial bone marrow niches are biologically distinct[28, 29], yet their relative sensitivity to rmTBI-induced hematopoietic alterations remains unknown. Given that the skull absorbs and transmits external mechanical forces during head trauma, it is plausible that calvarial bone marrow responds uniquely to rmTBI compared to distal marrow sites.

Here, we investigated the effects of rmTBI on calvarial and femoral hematopoiesis using a modified weight-drop model that delivers acceleration–deceleration and rotational forces to the brain[30]. We demonstrate that repeated sub-concussive head impacts induce successive waves of stem/progenitor cell proliferation resulting in early replicative senescence in both bone marrow compartments, accompanied by leukopenia and innate immune dysfunction. Moreover, secretory factors released from the rmTBI-exposed calvaria bone marrow compartment exhibit neurometabolic dysfunction and immunomodulatory effects on microglia. These immune alterations coincide with decreased skull bone mineral density, persistent neurological deficits, and regional microglial activation, identifying calvarial bone marrow as a previously underappreciated mediator linking repeated head trauma to chronic neuroimmune dysfunction.

## Methods

### Animals

Wildtype (WT) C57BL/6 male mice (8 weeks-old) were obtained from The Jackson Laboratory (Bar Harbor, ME, USA). Mice were group-housed in individually ventilated cage (IVC) racks with ad libitum access to standard chow and water. Housing conditions were maintained at a temperature of 21–23°C with a 12:12-hour light:dark cycle. All animal procedures were conducted in accordance with institutional guidelines at an AAALAC-accredited facility and were approved by the Animal Welfare Committee at the University of Texas Health Science Center at Houston (UTHSC-Houston), TX, USA.

### Repeated Mild Traumatic Brain Injury Model

To investigate the immunological and physiological consequences of rmTBI, a modified weight-drop model of closed-head injury was utilized, designed to recapitulate the mechanical forces commonly experienced by athletes in contact sports, including linear acceleration, deceleration, and rotational forces. Previous studies have reported that high school football players experience up to 650 impacts per season, while collegiate and professional players may sustain up to 1,400 impacts per season[31, 32]. Young adult male C57BL/6J mice (aged 8 weeks at the start of the experiment) were randomly assigned to rmTBI or sham control groups. Mice in the rmTBI group received a total of 48 head impacts over a 16-week period, with three impacts administered once weekly at one-hour intervals. This protocol was adapted from established models of repetitive TBI with long-term endpoints[30, 33, 34].

A standardized, commercially available weight-drop device (Conduct Science, Cat#: CS-WDTBIM) was used. A 90 g brass weight (2.5 cm diameter) was dropped from a height of 100 cm through a vertically aligned guide tube to ensure consistent impact. To minimize the risk of skull fracture and maximize consistency, the weight was fitted with a 4 mm-thick silicone gel padding (Dr. Scholl’s®). The impact was delivered to the vertex of the skull along the sagittal suture line, producing a glancing blow rather than a direct impact.

Prior to impact, mice were anesthetized using 4% isoflurane in an induction chamber. Anesthesia was confirmed by monitoring respiratory rate and the absence of the toe-pinch reflex. Anesthetized mice were positioned on their belly on a taut, single-ply low-lint laboratory wipe (Kimwipes®, Cat#: S-8116), suspended 11.5 cm above a polyurethane foam pad (38 cm × 18 cm × 25 cm). After initial impact, the mice passed through the wipe, rotated away from the weight impact point, and landed on the foam padding. Sham control animals were anesthetized for an equivalent duration without receiving an impact. Following each impact, mice were placed supine on a flat surface, and the latency to righting (i.e., turn over onto all four limbs) was recorded. Righting reflex time served as a proxy for loss of consciousness[35]. Mice were returned to their home cages between impacts, with a one-hour recovery period between each of the three weekly impacts.

Body weights were recorded weekly. To monitor for potential skull fractures or other structural injuries, micro-computed tomography (MicroCT) imaging was performed every four weeks. Scans were evaluated by institutional veterinary staff and an animal welfare administrator. Mice were monitored for three consecutive days following each weekly impact session using a study-specific ethogram developed for this experiment. Animals receiving a weekly ethogram score greater than six were flagged for clinical assessment by veterinary staff and were excluded from further participation in the study if warranted. Young adult male mice were selected to reflect the primary human population at risk for repetitive mild TBI-young adult males participating in contact sports. The selected closed-head modified weight-drop protocol is widely used and validated in preclinical TBI research[36, 37].

### Bromodeoxyuridine Administration

To measure *in vivo* Bromodeoxyuridine (BrdU) incorporation, mice were administered a dose of 50 mg/kg via i.p. injection (200 µL volume in PBS) at twelve hours prior to sacrifice. Body weights were recorded for each mouse and the appropriate dosages were administered.

### Behavioral Testing

Behavioral testing was performed over six consecutive days at three time points: baseline (−14 days), week 8, and week 16. Motor function was assessed using DigiGait, open field testing, and grip strength, while cognitive function was evaluated using the Novel Object Recognition Test (NORT) and fear conditioning. The order of behavioral assays was held constant across all testing periods to ensure consistency. Baseline and week 16 time points included the full behavioral battery to assess longitudinal changes, whereas week 8 testing was limited to grip strength to minimize habituation and accommodate ongoing repeated injury exposures. Fear conditioning was conducted only at week 16 to avoid potential confounding effects associated with repeated testing.

#### Open Field Test (OFT)

The OFT was used to assess spontaneous locomotor activity and anxiety-like behavior. Mice were placed in the center of an open-field chamber, and total distance traveled over a 20-minute session was quantified using the EthoVisionXT tracking system. Anxiety-like behavior was inferred based on the proportion of time spent in the periphery versus the center of the arena, with greater peripheral time indicative of increased anxiety.

#### Novel Object Recognition Test (NORT)

The NORT was used to evaluate recognition memory. Mice were habituated to the testing arena during the OFT. Twenty-four hours later, they were exposed to two identical objects for a total of 20 minutes, with exploration behavior recorded during those 20 minutes. On the same day, one hour later, one familiar object was replaced with a novel object, and exploration times were recorded. Time spent exploring the novel object and time spent exploring the familiar object were recorded. All data was quantified utilizing the EthoVisionXT tracking system software.

#### DigiGait

Motor coordination and gait were assessed using the DigiGait apparatus (Mouse Specifics, Inc.). Mice were placed on a transparent motorized treadmill that was gradually accelerated to a target speed of 15 cm/s, with younger mice tested at higher speeds when appropriate. DigiGait software was used to acquire and analyze stride length, swing time, ataxia, and additional gait parameters associated with white matter integrity.

#### Grip Strength

Grip strength was assessed using the BioSeb Grip Strength Meter. Mice were grasped by the base of the tail and lowered toward the grid apparatus, allowing both fore- and hindlimbs to attach. Mice were gently pulled back horizontally by the tail to elicit grip force. This procedure was repeated 10 times to obtain consistent measurements for both forelimb and hindlimb strength. Mouse weight was also recorded for normalization.

#### Whole-Body Plethysmography

Respiratory function was measured using the Whole-Body Plethysmography system (Prime Bioscience). Mice were singly housed in chambers for up to 2 hours: 1 hour for acclimation and 1 hour for recording. Parameters measured included respiratory rate (frequency), tidal volume, minute ventilation, and apneas.

#### Fear Conditioning

Contextual memory was assessed using a fear conditioning paradigm. On day 1, mice underwent habituation followed by foot-shock training in a rectangular conditioning chamber. On day 2, contextual memory was evaluated in the same chamber in the absence of shocks. Freezing behavior during the first minute of both training and testing sessions was quantified using EthoVision XT software, with reduced freezing during the testing session interpreted as impaired contextual memory.

### Tissue Collection

Two weeks after the final impact, mice were euthanized and tissues collected. Animals were weighed immediately prior to euthanasia, deeply anesthetized with isoflurane, and euthanized by exsanguination, with death confirmed by the absence of a pedal reflex. Blood was collected via external cardiac puncture using a 1 mL 25G tuberculin syringe (BD Biosciences, Cat# 309626) pre-coated with 0.5 M EDTA (pH 8.0; Invitrogen, Cat# 15575-038). Of the collected blood, 200 µL was reserved for flow cytometry, and the remaining volume was centrifuged at 12,000 rpm for 15 min at 4 °C to isolate plasma.

Following blood collection, mice underwent manual transcardial perfusion with 40 mL of ice-cold saline delivered at ∼10 mL/min using a 20 mL syringe (BD Biosciences, Cat# 302830). Brains, spleens, skulls, and femurs were harvested post-perfusion. Brains were hemisected, with tissue allocated for single-cell preparation and flow cytometry, flash-frozen for quantitative PCR, or fixed in 4% paraformaldehyde for 24 h before transfer to 70% ethanol for paraffin embedding and histopathological analysis.

### Skull Secretome Collection

Secretome samples were collected by submerging the calvaria (skullcap) in 2 ml of PBS for 2 hours in a 37C water bath. After 2 hours, the secretome was collected after removing the bone from the PBS and centrifuging the remaining liquid at 500g for 5 minutes at 4C. The supernatant was collected as the secretome sample and frozen at –80C until use.

### Multiplex Cytokine Enzyme-Linked Immunosorbent Assay

Levels of pro-inflammatory cytokine markers in the skull secretome were analyzed using a 23 - plex cytokine assay (Bio-Rad, #M60009RDPD). The assay was performed according to the manufacturer’s protocol, without prior dilution of the skull secretome samples. The samples were thawed on ice then loaded onto the plate per the manufacturer’s instructions, and incubation with detection antibodies and SA-PE was performed. The 96-well plate was read by the Bio-Plex 200 Suspension Array system, and the data was analyzed and acquired using the accompanying Bio-Plex Manager. Data are presented as pg/ml.

### Flow Cytometry

Whole blood was isolated by external cardiac puncture using EDTA-coated needles. Mice were then perfused with 40 ml of cold saline. 200 µl of blood was lysed for 10 min on ice using red blood cell lysis buffer (Biolegend, Cat# 420301) and then washed with FACS buffer. This was repeated up to three times until sufficient lysis was achieved. Bone marrow was harvested from the left femur by flushing with Roswell Park Memorial Institute (RPMI) (Lonza Group, Basel, Switzerland) medium using hydrostatic pressure. The calvaria (top portion of the skull) was excised using dissection scissors, and placed on ice in a 1.5 mL Eppendorf microcentrifuge tube containing 700 µL RPMI. After calvaria were collected, they were finely minced using scissors, and then passed through a 70-µm filter to remove debris. Brains were halved along the interhemispheric fissure to isolate one brain hemisphere. The olfactory bulb and cerebellum were then removed, and tissue was placed separately in complete Roswell Park Memorial Institute (RPMI) 1640 (Invitrogen, Cat# 22400105) medium and mechanically and enzymatically digested in collagenase/dispase (1 mg/ml, Roche Diagnostics, Cat# 10269638001), papain (5 U/ml, Worthington Biochemical, Cat# LS003119), 0.5 M EDTA (1:1000, Invitrogen, Cat# 15575020), and DNAse I (10 mg/ml, Roche Diagnostics, Cat# 10104159001) for 1 h at 37 °C on a shaking incubator (200 rpm). The cell suspension was washed twice with RPMI, filtered through a 70-µm filter, and RPMI was added to a final volume of 5 ml/hemisphere and kept on ice. Cells were then transferred into FACS tubes and washed with FACS buffer. Blood, bone marrow, and brain cells were then incubated with TruStain FcX Block (Biolegend, Cat# 101320), for 10 min on ice, and stained for the following surface antigens: CD45 -eF450 (eBioscience, Cat# 48–0451 − 82), CD11b-APCeF780 (eBioscience, Cat# 47011282), CD11c-PECy7 (Biolegend, Cat# 117318), MHCII-AF700 or PerCPCy5.5 (Biolegend, Cat# 107622/ 107626), Ly6C-AF700 or APC (Biolegend, Cat# 128024/128016), and Ly6G-PE or PerCPCy5.5 (Biolegend, Cat# 127607/127616). The fixable viability dye Zombie Aqua was used for live/dead discrimination (Biolegend, Cat# 423102). Cells were then washed in FACS buffer, fixed in 2% paraformaldehyde for 10 min, and washed once more prior to adding 500 µl FACS buffer. Intracellular staining for Ki67-PECy7 (Biolegend, Cat# 652426) and PCNA-AF647 (Biolegend, Cat# 307912) was performed using Cytofix/Cytoperm Fixation/Permeabilization Kit (BD Biosciences, Cat# 554714) according to manufacturer’s instructions and as described previously[38]. Cytokine staining for TNF-PECy7 (Biolegend, Cat# 506324) IL-6-APC (Biolegend, Cat# 504508) was performed after 3-hour incubation with Brefeldin A (Biolegend, Cat# 420601) at 37 °C followed by fixation/permeabilization. For phagocytic activity, cells were incubated with pHrodo™ Red Zymosan BioParticles (Invitrogen, Cat# P35364), according to the manufacturer’s instructions. Antigen presentation and T cell receptor (TCR) activation were measured by surface expression of MHCII and CD69-APC (Biolegend, Cat# 104514), respectively. For ROS detection, cells were incubated with DHR123 (5 mM; Life Technologies/Invitrogen, Cat# D23806), a cell permeable fluorogenic probe that reacts with hydrogen peroxide and superoxide. Cells were loaded for 20 min at 37 °C, washed three times with FACS buffer (without NaAz), and then stained for surface markers including viability stain.

To measure MitoSpy Red CMXRos (Biolegend, Cat# 42480) fluorescence intensity in secretome-stimulated samples: brain cells were washed once with PBS then incubated with 200ul of either PBS, sham skull secretome, or rmTBI skull secretome for 2 hours in a 37 °C water bath. Cells were washed twice with supplemented RPMI, then incubated with 200ul of MitoSpy Red CMXRos (Biolegend, Cat# 42480) at a concentration of 1:8000 in supplemented RPMI for 25 minutes at 37 °C. Cells were washed with FACS buffer, then stained with TruStain FcX Block (Biolegend, Cat# 101320), for 10 min on ice, followed by staining for viability and surface markers as explained above.

Data were acquired on a 5-laser CytoFlex LX cytometer (Beckman-Coulter) using Cytek software (Cytek Biosciences) and analyzed using FlowJo (Treestar Inc.). At least 5–10 million events were collected for each sample. CountBright Absolute Counting Beads (ThermoFisher, Cat# C36950) were used to estimate cell counts per the manufacturer’s instructions. Data were expressed as counts/hemisphere (brain) or counts/μL (blood, bone marrow). Leukocytes were first gated using a splenocyte reference (SSC-A vs. FSC-A). Singlets were gated (FSC-H vs. FSC-W), and live cells were gated based on Zombie dye exclusion (SSC-A vs. Zombie Aqua-Bv510). Bone marrow LSK + cells were identified as lineage^−^c-Kit^+^Sca1^+^ (Biolegend, Cat# 133307, 405226, 105837, and 108111). Monocytes and neutrophils were identified as CD45^+^CD11b^+^Ly6C^+^Ly6G^-^ and CD45^+^CD11b^+^Ly6C^+^Ly6G^+^, respectively (**Supplemental Figure 1**)[39]. Resident microglia were identified as the CD45^int^CD11b^+^Ly6C^−^ population, whereas peripheral leukocytes were identified as CD45 ^hi^CD11b^+^ myeloid cells or CD45^hi^CD11b^−^ lymphocytes. Cell type–matched fluorescence minus one (FMO) controls were used to determine the positivity of each antibody.

### Mass Spectrometry-based Proteomics

The secretome samples collected in PBS buffer from skull tissue were further diluted in 50mM final ammonium bicarbonate. The solution was snap frozen in liquid nitrogen and boiled on a heat block at 950C for 5min. The lysate was allowed to cool at room temperature and protein concentration measured using Bradford assay. The protein digestion was carried out using trypsin and LysC enzyme mixture at 37C overnight. The enzyme reaction was neutralized by adding 1% final formic acid. The peptide sample was desalted on C18 stage tips and measured using the Pierce™ Quantitative Colorimetric Peptide Assay (Thermo Scientific 23275). The peptides were concentrated in a speed vac and dissolved in MS loading buffer (0.015% DDM prepared in 0.1% formic acid). The peptide samples were measured on a nanoElute2 liquid chromatography system (Bruker Daltonics, Germany) coupled online to timsTOF Ultra2 mass spectrometer (Bruker Daltonics, Germany) via a CaptiveSpray nano-electrospray ion source. The data acquisition was performed in diaPASEF mode. The peptide samples (50ng) were directly loaded onto a 25cm x 75µm Aurora Ultimate column (IonOpticks) and separated using a 30-min total run time. Eluting peptides were measured using the default ‘low-input amount dia-PASEF’ method on tims Control (v6.0). The dia-PASEF window scheme included m/z range (400 to 1000m/z) with a mobility range (0.64 to 1.37 1/K0). The diaPASEF raw data was processed using DIA-NN (v 2.1.0) in a library-free search mode. The fasta file containing 17,228 mouse proteins was downloaded from UniprotKB (downloaded 2024_11_21). The precursor ions were generated from FASTA in silico digest with deep learning-based spectra, retention times (RTs), and ion mobilities (IMs) prediction. The proteolytic enzyme Trypsin/P was selected, allowing for up to one missed cleavage. Variable modification of oxidation (M) and protein N-term acetylation was allowed. Parameter specifics include peptide lengths ranging from 7–30, precursor charges from 1–4, precursor m/z from 300–1800, and fragment ion m/z from 200–1800. The MBR was enabled and precursor identification was set at 1%FDR. The gene product inference and quantification was done with label-free iBAQ approach using ‘gpGrouper’ algorithm[40]. The iBAQ protein expression was median normalized and log-transformed for further statistical analysis.

### qPCR

Total RNA was extracted from mouse brain hemisphere homogenates using the RNeasy Mini Kit (QIAGEN, Cat# 74104), following the manufacturer’s instructions. For each sample, 500 ng of RNA was reverse transcribed into complementary DNA (cDNA) using the RevertAid First Strand cDNA Synthesis Kit (Thermo Scientific, Cat# K1621). Quantitative real-time PCR was performed using either the Bio-Rad CFX384 or CFX96 Real-Time PCR Detection Systems. Each 20 µL PCR reaction contained 1 µL of cDNA template, 10 µL of TaqMan Fast Advanced Master Mix (Thermo Fisher Scientific, USA), and 1 µL of gene-specific TaqMan probe (Thermo Fisher, Cat# 4331182), with the remaining volume adjusted using nuclease-free water. Relative mRNA expression levels were calculated using the 2^−ΔΔCt method. Target gene expression for *p53* (TaqMan, Catalog # Mm03976700_m1), *p27* (TaqMan, Catalog # Mm00438168_m1), and *p21* (TaqMan, Catalog # Mm00432448_m1) was normalized to *Gapdh* (TaqMan, Catalog # Mm99999915_g1) as the internal control.

### Measurement of Relative Telomere Length

Flash-frozen skull and femur cells were collected from mice, and DNA was extracted using the QIAamp DNA Mini Kit (Qiagen, Cat. No. 51306) according to the manufacturer’s instructions. DNA concentration and purity were assessed using a NanoDrop microvolume spectrophotometer (Thermo Scientific, Cat. No. 13-400-518). Samples were diluted to a final concentration of 60 ng/µL using DNA/RNA-free water. Quantitative PCR (qPCR) reactions were prepared using 12.5 µL SYBR Green Supermix (Bio-Rad, Cat. No. 1725271), 0.7 µL each of telomeric forward and reverse primers, 0.5 µL each of albumin forward and reverse primers, and template DNA in a final reaction volume adjusted with nuclease-free water. Reactions were run in technical triplicates on a CFX Opus 384 Real-Time PCR System (Bio-Rad, Cat. No. 12011452). This procedure was based on the protocol established in Jiao et al. 2012. The primer sequences used were as follows:

### Telomeric forward primer

5′-ATACCAAGGTTTGGGTTTGGGTTTGGGTTTGGGTTCATGG-3′

### Telomeric reverse primer

5′-GAGGCAATATCCCTATCCCTATCCCTATCCCTATCCCTAACC-3′

### Albumin forward primer

5′-CGGCGGCGGGCGGCGCGGGCTGGGCGGAAACGCTGCGCAGAATCCTTG-3′

### Albumin reverse primer

5′-GCCCGGCCCGCCGCGCCCGTCCCGCCGCTGAAAAGTACGGTCGCCTG-3′

### Dual-Energy X-ray Absorptiometry (DEXA)

Following euthanasia, skulls were excised by carefully cutting along the lateral aspects of the skull. The skull cap was removed and immediately placed in RPMI medium (Lonza Group, Basel, Switzerland) before being transported to the DEXA InAlyzer2, Model S apparatus for analysis. To ensure consistency across samples, each skull cap was trimmed to achieve a comparable surface area. The trimmed skull tissue was positioned within the scanning field, and user-defined regions of interest (ROIs) were selected for each sample. All ROIs were consistently drawn to encompass the entire surface area of the excised skull tissue. A specialized “excised tissue” scan protocol was executed, and bone mineral density data were collected for each sample.

### MicroCT

Every 4 weeks, to confirm no skull fracturing was taking place, MicroCT imaging was performed on a Bruker SkyScan 1276 CMOS imaging system equipped with an isoflurane anesthesia vaporizer and nose cone. Heat support was provided to the mouse during imaging. The X-ray source was 60 kV and 200 uA and passed through a 0.5 mm Aluminum filter. Projection images were obtained with 4×4 camera binning at 0.8 degree rotational interval (315 msec exposure) for final pixel size of 37.3 microns. Total estimated radiation exposure was 500 mGy per imaging session. Subsequently, 3D image reconstruction was performed with NRecon (Bruker) and analysis was performed with DataViewer and CTvox (Bruker).

### Mitochondrial Bioenergetics

#### Preparation of Brain Slices

Naïve adult mice (n = 3; male, 3-4mo) were humanely euthanized, brains were rapidly removed (within 30 s of decapitation) and immersed in ice-cold (4–5 °C) artificial cerebrospinal fluid (aCSF; 120 mM NaCl, 3.5 mM KCl, 1.3 mM CaCl2, 1 mM MgCl2, 0.4 mM KH2PO4, 5 mM HEPES, and 10 mM D-glucose; pH 7.4) that had been oxygenated for 1 h using 95% O2:5% CO2. Coronal sections (200 μm) were prepared using a modified McIlwain tissue chopper (Ted Pella. Inc.; Redding, CA) with a chilled stage and blade, then transferred to a holding chamber containing continuously oxygenated aCSF at room temperature (∼ 23 °C).

#### Secretome Incubation

Secretome samples previously isolated from 5 TBI and 5 sham subjects were combined to provide adequate volume for *ex vivo* brain slice incubation and to eliminate the possibility of effects due to any one secretome sample. The secretome was added a 2:1 dilution of oxygenated aCSF to prevent hypoxia and/or nutrient deprivation and bath applied at a 30% final concentration to brain slices. This was repeated 3 times, 60 minutes apart to constantly ensure oxygenation (a protocol previously used to assess mitochondrial changes after chemical LTP[41]. TBI and sham secretome effects on healthy brain tissue were compared to samples incubated with aCSF alone to control for any influence of the incubation protocol on mitochondrial function.

#### Tissue Punches and Respiration Measurements

Brain sections were individually transferred to a biopsy chamber containing fresh oxygenated aCSF. A stainless steel WellTech Rapid-Core biopsy punch needle (500 μm diameter; World Precision Instruments; Sarasota, FL) was used to excise the tissue punches. Tissue punches were taken from each anatomical location using four consecutive coronal sections (*i.e.*, a total of 4 punches for each anatomical structure) using the same biopsy punch needle. Punches were ejected directly into an XF96 Pro Cell Culture Microplate (101085-004; Agilent Technologies, Santa Rosa, CA) based on a predetermined plate layout. Each well contained 180 μL room temperature assay media (aCSF supplemented with 0.6 mM pyruvate and 4 mg/mL lyophilized BSA). After loading all biopsy samples, each well was visually inspected to ensure that the punch was submerged and centered at the bottom. The XF96 Pro Cell Culture Microplate was then incubated at 37 °C for approximately 30 min. During this incubation period, 10 × concentration of assay drugs (prepared in aCSF) were loaded into their respective injection ports of a hydrated (overnight in distilled water, exchanged for XF Calibrant solution 3 h prior to assay initiation) Seahorse XF96 Pro Extracellular Flux Assay sensor cartridge. The sensor cartridge containing the study drugs was then inserted into the analyzer for calibration. Once the analyzer was calibrated, the calibration plate was replaced by the microplate containing the tissue punches and the assay protocol initiated. Assay drugs were prepared at 10 × working concentrations in oxygenated aCSF (pH 7.4) and delivered sequentially to achieve final concentrations of: port A: Oligomycin (25 μg/mL); port B: FCCP + pyruvate (75 μM and 7.5 mM, respectively); and port C: Antimycin A + rotenone (10 μM and 5 μM, respectively). The dose of each of these reagents was based on optimization experiments to achieve optimum drug effect[42]. The duration of sampling time (for calculating the oxygen consumption rate or OCR) for each condition was determined to allow the effect of each drug to reach a steady state. Wells with low basal activity (< 20 pmol/min OCR) and/or failing to respond to FCCP/pyruvate were excluded from analysis. Tissue respiration results were normalized following the procedures recommended by the Seahorse XF analyzer manufacturer for 3-D samples (validated in[41]).

### Statistical Analysis

All statistical analyses were conducted using GraphPad Prism 7 (GraphPad Software, San Diego, CA, USA). Group comparisons were performed using an unpaired parametric Student’s *t*-test for two-group analyses, and two-way repeated measures ANOVA was used to assess interaction effects over time. Where applicable, main effects were tested, followed by post hoc analyses with *p*-values adjusted for multiple comparisons using Sidak’s correction. Spearman’s rank correlation coefficient was used to assess statistical associations between average behavioral variables and levels of flow cytometry and immunohistochemical markers within each experimental condition. Outliers were identified post hoc using Grubb’s test and excluded from analysis when statistically justified. Statistical significance was defined as *p* < 0.05. In all figures, significance is denoted as follows: * *p* < 0.05, ** *p* < 0.01, *** *p* < 0.001, and **** *p* < 0.0001.

## Results

### Mild impairments in cognitive, motor, and respiratory function are evident in a 16-week rmTBI model with inter-session recovery

To begin our investigation into the cumulative effects of repeated head injury on hematopoietic bone marrow responses, mice were subjected to a rmTBI paradigm consisting of once-weekly injury sessions for 16 consecutive weeks (**Figure 1A**). Each injury session comprised three consecutive mild weight-drop impacts delivered at 1-h intervals. This schedule was designed to model repeated sub-concussive head impacts while allowing inter-session recovery. Behavioral testing was conducted throughout our 16-weeks of rmTBI to examine neurological outcomes. No change in righting reflex times was noted until after 11 -weeks of rmTBI, compared to sham controls (**Figure 1B**). These results, although variable, suggest that the compounding effects of rmTBI may progressively increase injury severity associated with prolonged loss of consciousness. Body weight loss was largely unperturbed after rmTBI (**Figure 1C**), highlighting the mild nature of the injury. Moreover, no changes in spontaneous locomotor activity or body speed were seen in the open field test (**Figures 1D-F**). Significant reductions in grip strength force were found in rmTBI, but not sham mice, over time (**Figure 1G**). Next, to measure recognition memory, we performed the novel object recognition test after 16-weeks of rmTBI. rmTBI mice made significantly fewer entries into the novel object zone, and spent less time interacting with the novel object compared to sham controls (**Figures 1H-J**), indicating greater cognitive impairment. Gait analysis using the DigiGait system, revealed significantly slower hindlimb movements and increased forelimb ataxia after 16-weeks of rmTBI compared to sham controls (**Figures 1K-O**). Finally, our plethysmography results after 16-weeks of rmTBI showed no change in heart beats per minute or minute ventilation between groups (**Figures 1P-R**). However, significant increases in tidal volume and apneas per minute were found ( **Figures 1S-T**), suggesting that rmTBI mice experience more frequent pauses in breathing (apneas), but compensate by taking deeper breaths (increased tidal volume) to maintain normal overall ventilation. Altogether, our comprehensive battery of behavioral tests suggests our rmTBI model is indeed mild, and is associated with chronic and diverse neurological impairments.

**Figure 1.**
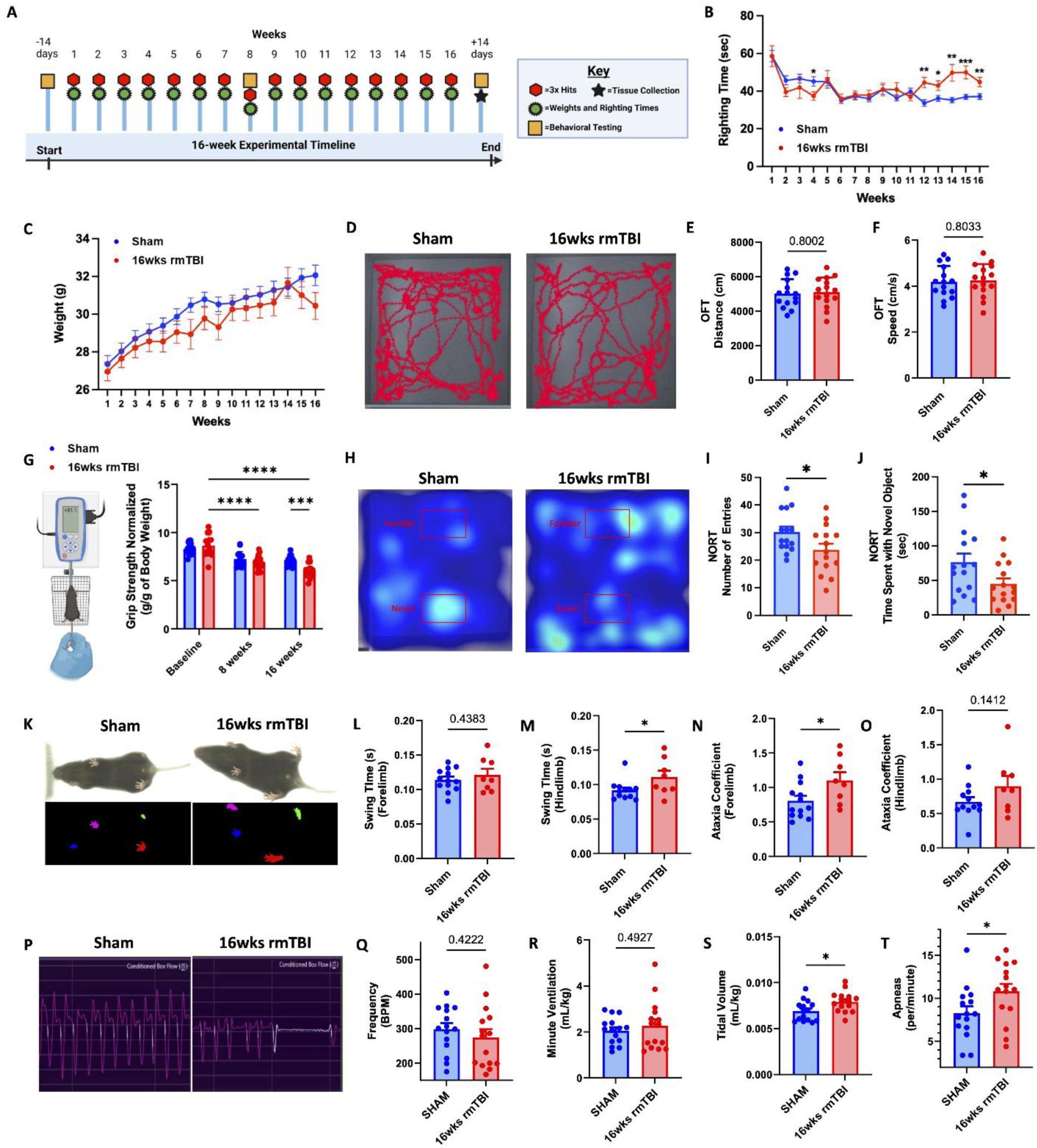
Mild cognitive, motor, and respiratory dysfunction in a 16-week rmTBI model with inter-session recovery. Diagram of the behavioral testing timeline employed to determine neurological outcome throughout 16-weeks of rmTBI (**A**). Righting reflex times and body weights were recorded at repeated timepoints for sham and rmTBI groups (**B** and **C**, respectively). Representative movement trace plots (**D**) show no significant change in total distanced traveled (**E**) or body speed (**F**) in an open field. Grip strength was measured at 8- and 16-weeks after rmTBI (**G**). Image provided by *Biorender.com*. Representative spatial heatmaps of object exploration in the novel object recognition test are shown (**H**) and number of entries (**I**) and time spent with the novel object (**J**) were quantified after 16-weeks of rmTBI. Representative gait dynamics traces from DigiGait analysis are shown (**K**) and fore- and hind-limb swing times (**L-M**) and ataxia coefficients (**N-O**) were quantified. Representative respiratory waveform traces from plethysmography testing are shown (**P**). Heart rate frequencies (**Q**), minute ventilation (**R**), tidal volume (**S**), and apneas (**T**) for sham and rmTBI groups were quantified. Data were analyzed using Student’s T-test (**E-F**, **I-T**) or repeated measures ANOVA with Tukey’s post-hoc test for multiple comparisons (**B-C**, **G**). \*\*\*\**P* < 0.0001, \*\*\**p* < 0.001, and \**p* < 0.05. rmTBI: repeated mild traumatic brain injury, BPM beats per minute, cm centimeter, g gram, kg kilogram, ml milliliter, s/sec seconds, wk weeks.

### rmTBI induces acute stem/progenitor cell proliferation and myeloid output in femoral and calvarial bone marrow

We next assessed the acute responsiveness of femoral and calvarial bone marrow stem/progenitor (LSK⁺) cells to rmTBI 24 h after either 1 or 2 weeks of injury (three consecutive impacts/week; **Figure 2A**). In the femur, rmTBI did not alter total LSK cell numbers at either time point (**Figure 2B**), but significantly increased the proportion of BrdU⁺ LSK cells following both 1 and 2 weeks of injury, indicating enhanced proliferation (**Figure 2C**). This proliferative response was accompanied by increased production of Ly6C⁺ monocytes, with no corresponding change in neutrophil output (**Figure 2D**).

**Figure 2.**
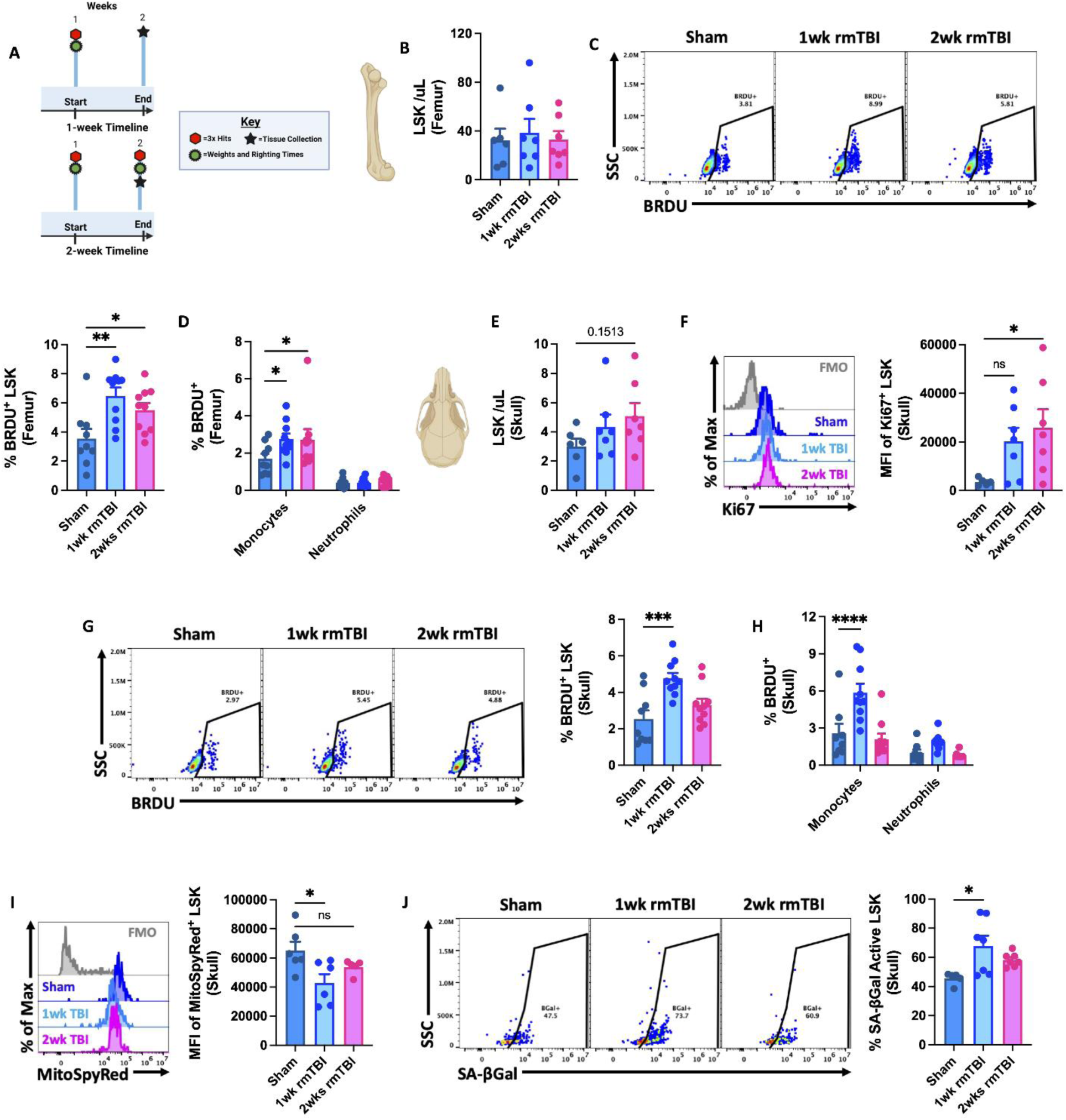
Comparison between local and distal bone marrow responses to 1- and 2-weeks of rmTBI. Diagram of the experimental timeline used to determine initial and subsequent bone marrow responses to 1- and 2-weeks of rmTBI (**A**). The concentration of LSK cells in the femur was quantified at both endpoints (**B**). The percentage of BrdU-positive LSK cells (**C**) and myeloid cells (**D**) in the femur is shown. The concentration of LSK cells in the calvaria of the skull is shown for both endpoints (**E**). Representative histograms depict the mean fluorescence intensity (MFI) of Ki67 protein expression in skull LSKs is elevated after rmTBI (**F**). The frequency of BrdU-positive skull LSKs (**G**) and myeloid cells (**H**) was quantified. Mean fluorescence intensity of MitoSpy Red CMXRos in skull LSKs was reduced after rmTBI (**I**). The percentage of senescence-associated beta-galactosidase active skull LSKs was significantly increased after the first of week of rmTBI (**J**). For all histograms, light gray = fluorescence minus one (FMO) control. Data were analyzed using one-way ANOVA (**B-C**, **E-G**, **I-J**) or two-way ANOVA analysis with Tukey’s test for multiple comparisons (**D**, **H**). \*\*\*\**P* < 0.0001, *** *p* < 0.001, \*\**p* < 0.01, and \**p* < 0.05. rmTBI: repeated mild traumatic brain injury, max maximum, SSC side scatter, µl microliter, wk weeks.

In the calvaria, rmTBI similarly did not change LSK cell numbers (**Figure 2E**). However, despite elevated Ki67 expression (**Figure 2F**), skull LSK cells exhibited reduced BrdU incorporation after the second week of injury, suggesting diminished proliferative capacity with repeated exposure (**Figure 2G**). Consistent with this finding, de novo monocyte production in the calvaria was observed only after the first week of head impacts (**Figure 2H**). Notably, rmTBI induced a significant reduction in mitochondrial membrane potential, as evidenced by MitoSpy Red CMXRos fluorescence intensity, in calvarial LSK cells during the first week (**Figure 2I**). This is indicative of impaired metabolic activity, and was accompanied by an acute increase in senescence-associated β-galactosidase activity (**Figure 2J**).

Collectively, these data demonstrate that rmTBI acutely stimulates LSK cell proliferation and monocyte production in both femoral and calvarial bone marrow. In contrast, skull-derived LSK cells rapidly develop metabolic dysfunction and senescence-associated features, which may underlie their attenuated responsiveness to subsequent repeated head impacts.

### Emergence of a senescent-like phenotype in skull bone marrow stem/progenitor cells after 8-weeks of rmTBI

After 8 weeks of rmTBI (three impacts/week; 24 total impacts), calvarial LSK cells failed to mount a proliferative response, as indicated by unchanged BrdU incorporation (**Figures 3A-B**). Although mitochondrial membrane potential showed a downward trend, this change did not reach statistical significance (**Figure 3C**). In contrast, senescence-associated β-galactosidase activity remained significantly elevated in calvarial LSK cells at this time point (**Figure 3D**), accompanied by increased expression of senescence-associated tumor suppressor genes as measured by qPCR (**Figure 3E**).

**Figure 3.**
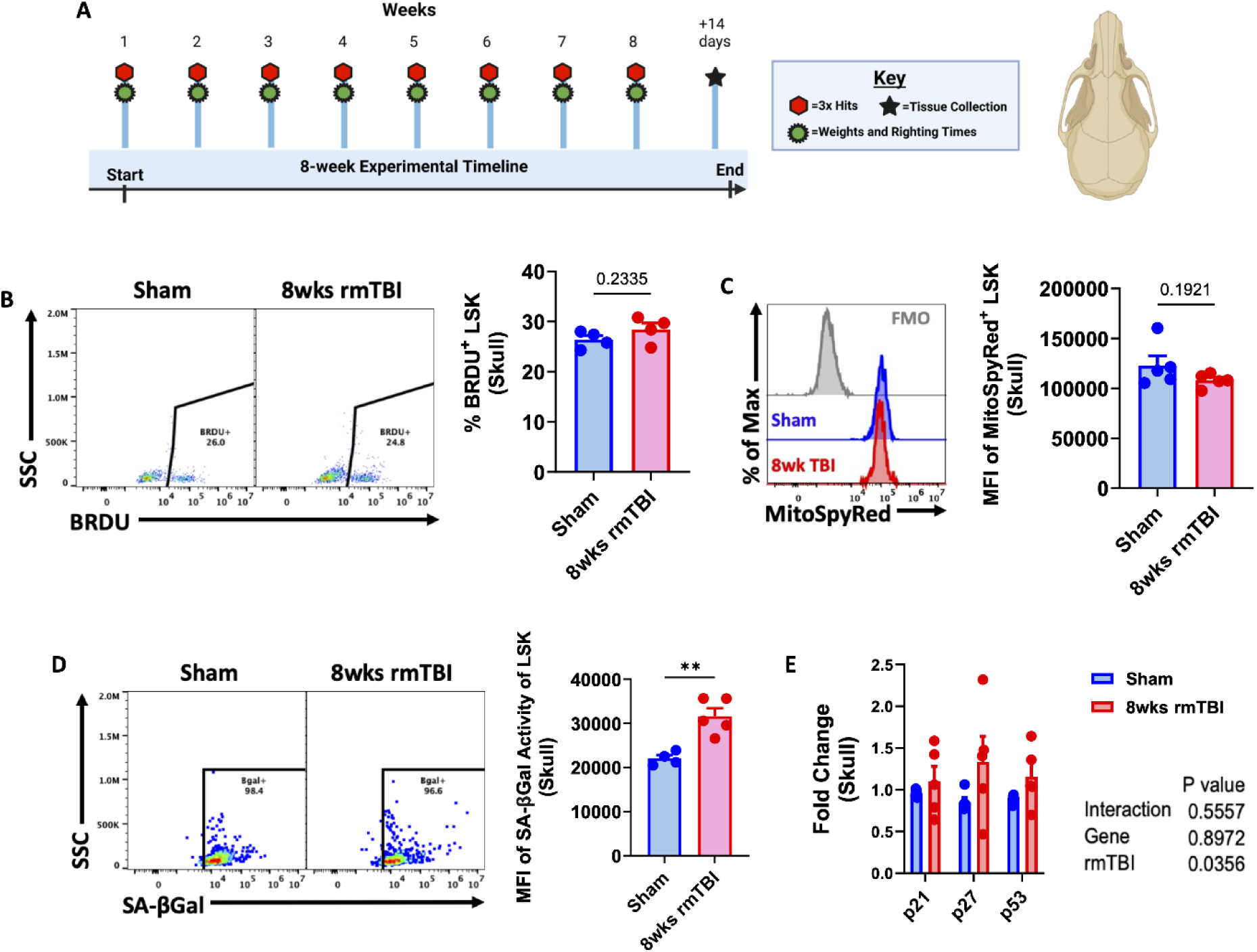
Altered skull bone marrow response after 8-weeks of rmTBI. Diagram of the experimental timeline used to determine skull bone marrow responses to 8 -weeks of rmTBI (**A**). The frequency of BrdU-positive skull LSKs is shown (**B**). Mean fluorescence intensity (MFI) of MitoSpy Red CMXRos in skull LSKs was unchanged after rmTBI (**C**). MFI of senescence-associated beta-galactosidase activity in skull LSKs was increased after rmTBI (**D**). Relative expression of the senescence-related tumor suppressor genes p21, p27, and p53 in skull bone marrow cells is shown (**E**). For all histograms, light gray = fluorescence minus one (FMO) control. Data were analyzed using Student’s T-test (**B-D**) or two-way ANOVA group analysis (**E**). \*\**P* < 0.01. rmTBI: repeated mild traumatic brain injury, max maximum, SSC side scatter, wk weeks.

Collectively, these findings indicate that rmTBI induces the emergence of a senescent-like phenotype in skull bone marrow stem/progenitor cells by 8 weeks, characterized by sustained senescence markers and impaired proliferative capacity.

### Bone marrow stem/progenitor cell deficiencies after 16-weeks of rmTBI

After 16 weeks of rmTBI (three impacts/week; 48 total impacts), followed by a 2 -week washout period, LSK stem/progenitor cells were significantly depleted in both the femur and calvaria (p = 0.02; **Figures 4A-B**). Consistent with this depletion, the proportion of BrdU⁺ LSK cells was markedly reduced in both bone marrow compartments (p = 0.004; **Figure 4C**), indicating diminished proliferative capacity. Two-way ANOVA revealed higher senescence-associated β-galactosidase activity in calvarial compared with femoral LSK cells (p = 0.006), as well as a significant increase following rmTBI (p = 0.04; **Figure 4D**). In parallel, analysis of leukocyte telomere length demonstrated significant telomere shortening in both femoral and calvarial bone marrow cells relative to sham controls (**Figures 4E-F**).

**Figure 4.**
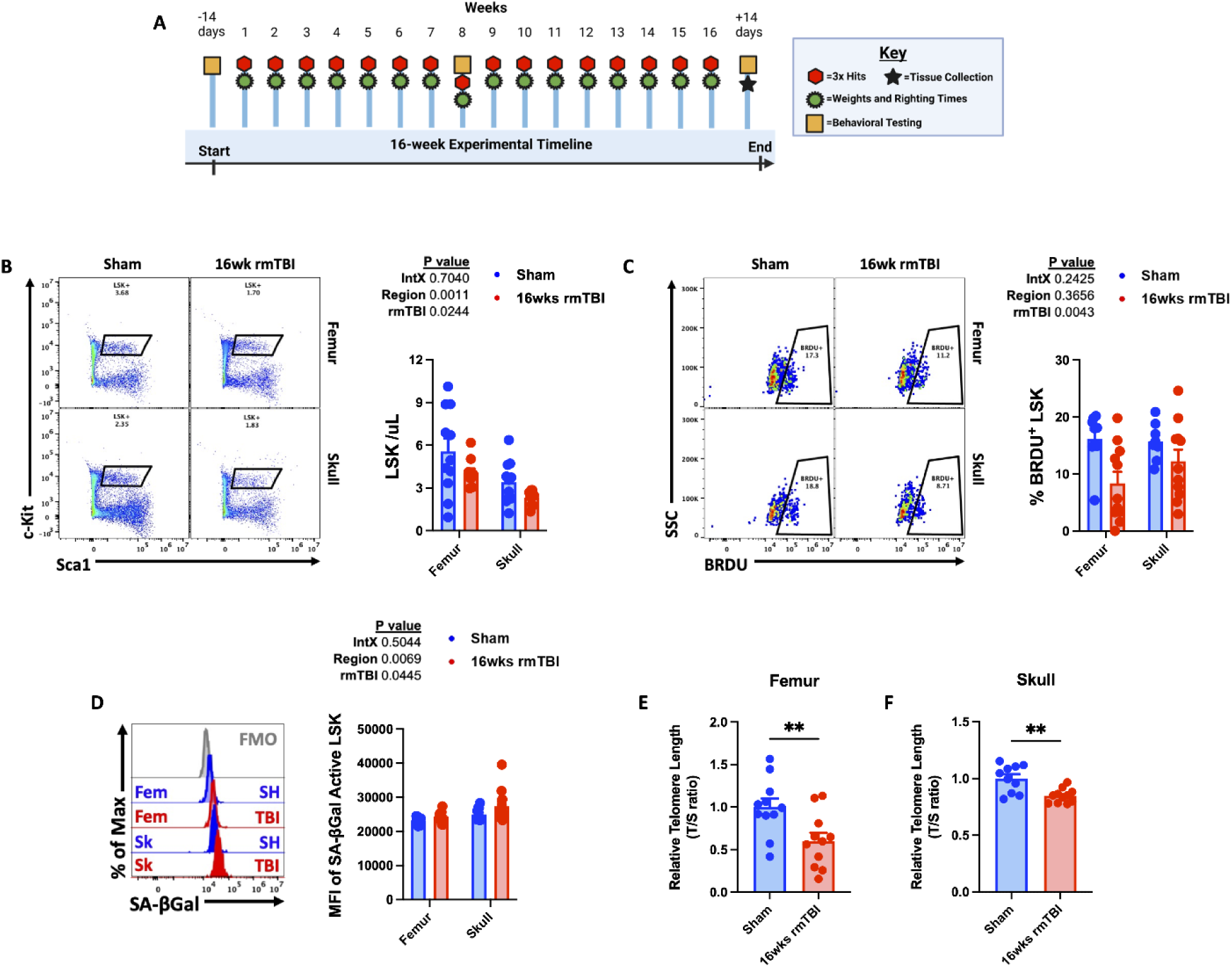
Evidence for replicative senescence in bone marrow after 16-weeks of rmTBI. Diagram of the experimental timeline used to determine skull bone marrow responses to 16-weeks of rmTBI (**A**). Representative dot plots depict frequency of LSK cells in the femur and skull after rmTBI (**B**). Representative dot plots show the percentage of BrdU-positive LSK cells after rmTBI (**C**). Mean fluorescence intensity (MFI) of senescence-associated beta-galactosidase activity in femur and skull LSK cells is shown (**D**). The relative telomere lengths of femur (**E**) and skull (**F**) bone marrow cells after 16-weeks of rmTBI. For all histograms, light gray = fluorescence minus one (FMO) control. Data were analyzed using two-way ANOVA group analysis (**B-D**) or Student’s T-test (**E-F**). \*\**P* < 0.01. rmTBI: repeated mild traumatic brain injury, intX interaction, max maximum, SSC side scatter, T/S telomere repeat copy number / single-copy gene number, µl microliter, wk weeks.

Together, these data indicate that chronic rmTBI leads to progressive functional impairment of bone marrow stem/progenitor cells, culminating in cellular depletion, reduced proliferative capacity, increased senescence-associated activity, and telomere attrition-hallmarks of injury-induced immune senescence.

### rmTBI-induces bone marrow hypocellularity, blood leukopenia, and innate immune dysregulation

To examine the long-term consequences of rmTBI on bone marrow hematopoiesis, we quantified leukocyte counts and *de novo* production. Significant decreases in skull-derived total CD45+, CD11b+, monocytes, neutrophils, and B cells were seen after 16 -weeks of rmTBI (**Figures 5A-E**). The percentage of BrdU-positive monocytes, neutrophils, and B cells were also significantly reduced in the skull and femur (**Figures 5F-K**), suggesting impaired production of hematopoietic cells. To confirm this finding, we quantified the number of leukocytes in blood. Significant corresponding reductions in circulating blood leukocytes were also seen (**Figures 6A-D**). Moreover, the relative level of reactive oxygen species (ROS) in blood leukocytes was significantly higher after rmTBI compared to sham controls (**Figures 6E-G**), suggesting increased oxidative stress. Next we assessed innate immune activation phenotypes across several peripheral lymphoid tissues. A non-significant (p=0.051) trend for lower thymus tissue weights was seen after rmTBI compared to sham control (**Figure 6H**). We also observed a non-significant (p=0.053) trend for lower CD4:CD8 T cell ratios in the spleen after rmTBI, and higher TNF production in CD8 T cells (**Figures 6I-J**). In the lungs, we found a decrease in zymosan engulfment by neutrophils, and in the mesenteric lymph nodes we found a significant down-regulation of MHCII-mediated antigen presentation that corresponded to a decrease in CD4 T cell TCR activation, as evidenced by lower proportions of CD69+ subsets (**Figures 6K-M**). These results indicate that the accumulated chronic stress of rmTBI may cause bone marrow failure, reducing the protective, reparative and regenerative capacity of the immune system to support healthy central nervous system function.

**Figure 5.**
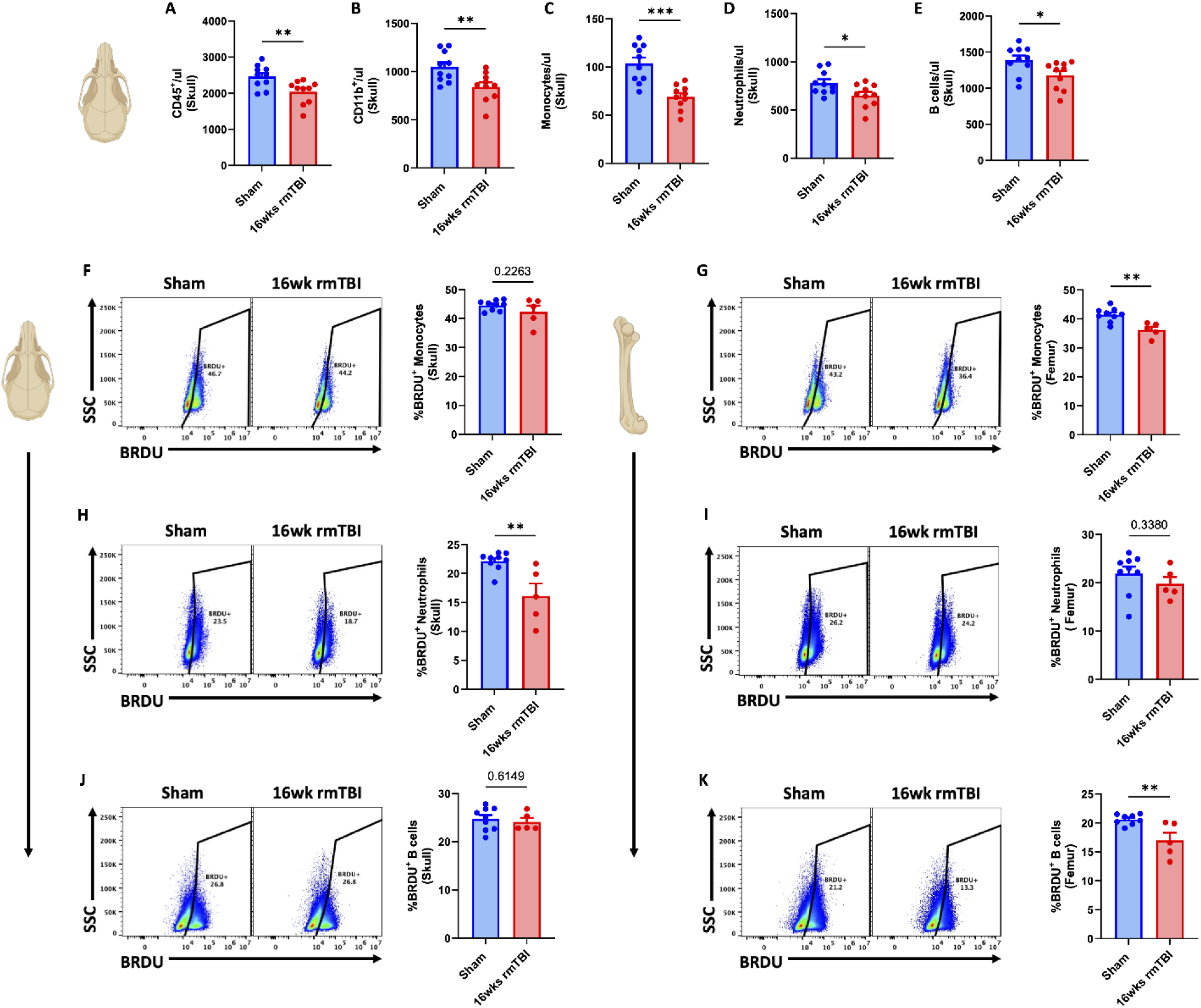
16-weeks of rmTBI causes reductions in bone marrow leukocyte production. Cell count concentrations were quantified for total CD45+ (**A**), total CD11b+ (**B**), monocytes (**C**), neutrophils (**D**), and B cells (**E**) in skull bone marrow after 16-weeks of rmTBI. Representative dot plots show the frequency of BrdU-positive monocytes (**F**), neutrophils (**H**), and B cells (**J**) in the skull. Representative dot plots show the frequency of BrdU-positive monocytes (**G**), neutrophils (**I**), and B cells (**K**) in the femur. Data were analyzed using Student’s T-test. \*\*\**P* < 0.001, \*\**p* < 0.01, and \**p* < 0.05. rmTBI: repeated mild traumatic brain injury, max maximum, MFI mean fluorescence intensity, SSC side scatter, µl microliter, wk weeks.

**Figure 6.**
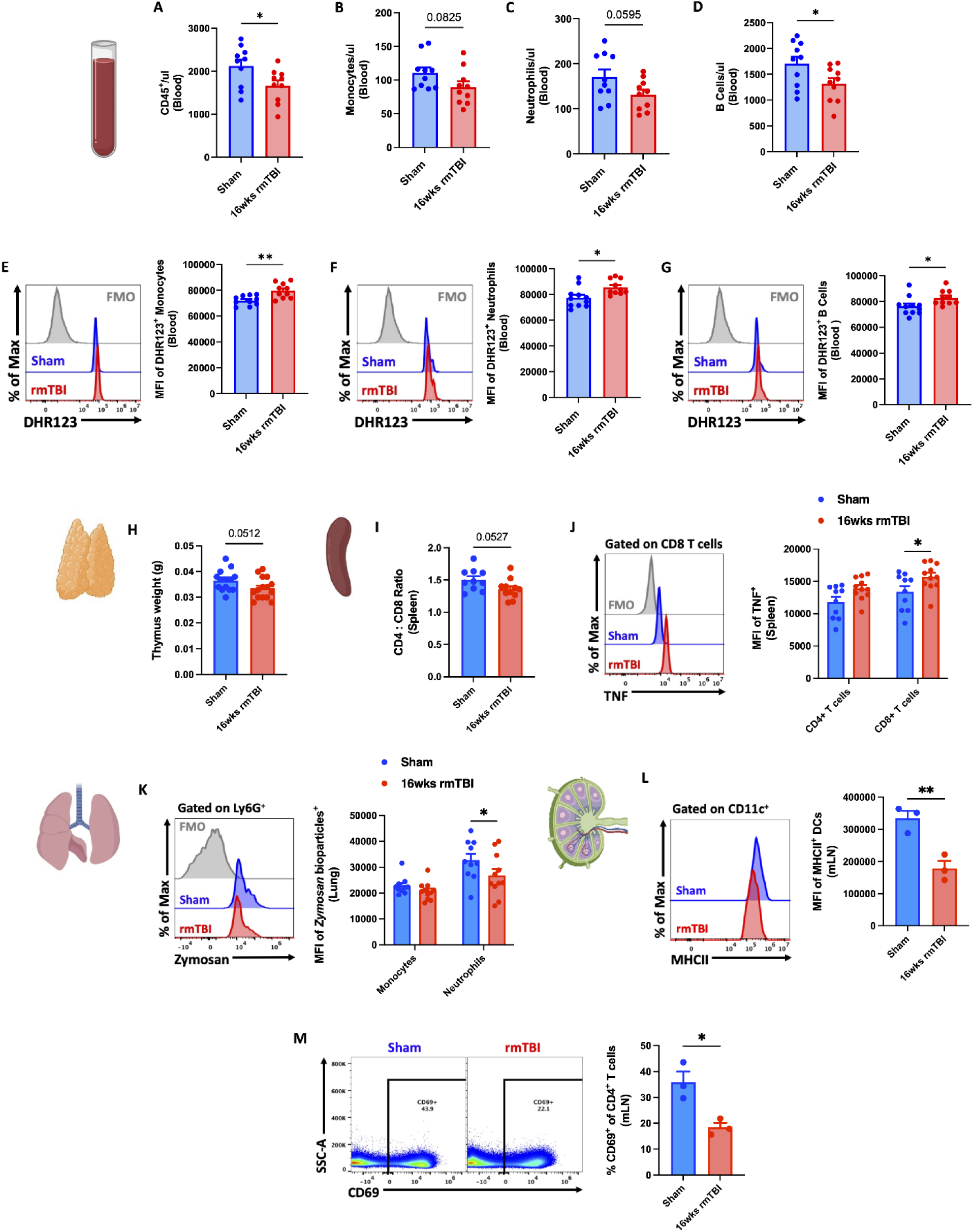
16-weeks of rmTBI causes leukopenia and innate immune dysfunction. Blood count concentrations are shown for total CD45 (**A**), monocytes (**B**), neutrophils (**C**), and B cells (**D**) after 16-weeks of rmTBI. Representative histograms depict the relative level of reactive oxygen species production in circulating blood monocytes (**E**), neutrophils (**F**), and B cells (**G**) as measured using dihydrorhodamine (DHR) 123 probe. Thymus weights (**H**) and CD4:CD8 ratios in the spleen (**I**) showed a non-significant decreasing trend after rmTBI. A representative histogram depicts the relative production of the pro-inflammatory Th1 cytokine, TNF, is shown for CD4 and CD8 T cell subsets in the spleen (**J**). Representative histograms show engulfment of pHrodo-conjugated zymosan bioparticles in lung-associated monocytes and neutrophils (**K**). Mean fluorescence intensity of MHCII was significantly decreased in mesenteric lymph node-associated dendritic cells (**L**). Representative dot plots show a significant decrease in the percentage of CD69-positive CD4 T cells in the mesenteric lymph nodes (**M**). For all histograms, light gray = fluorescence minus one (FMO) control. Data were analyzed using Student’s T-test and two-way ANOVA with Tukey’s post-hoc test for multiple comparisons. \*\**p* < 0.01, and \**p* < 0.05. rmTBI: repeated mild traumatic brain injury, DCs dendritic cells, mLN mesenteric lymph node, max maximum, MFI mean fluorescence intensity, SSC side scatter, µl microliter, wk weeks.

### rmTBI causes increased IL-6 secretion and decreased bone mineral density in the skull

Bone marrow failure resulting from dysregulated signals (e.g., cytokines, growth factors) can impair osteoblast differentiation and bone formation. We confirmed that our 16-week rmTBI model did not cause overt skull fracturing using microCT (**Figure 7A**). However, DEXA scan analysis revealed that bone mineral density was significantly decreased after 16-weeks of rmTBI in skull, but not femur (**Figures 7B-D**). To assess which cytokine and growth factor signals may be responsible for these changes, we assayed the calvarial secretome of sham and rmTBI mice using multiplex ELISA (**Figures 7E-J**). These data highlight a profound increase in IL-6 secretion (**Figure 7E**), and decrease in CCL3 (MIP-1α) concentrations (**Figure 7J**), after rmTBI. We then confirmed that IL-6 production was significantly increased in skull-, but not femur-, derived monocytes and neutrophils after 16-weeks of rmTBI, relative to sham controls (**Figures 7K-M**). Taken together, our discovery that IL-6 levels are significantly and chronically elevated after rmTBI are consistent with a senescence-associated secretory phenotype (SASP) and may consequently drive osteolytic activity and reduce bone density.

**Figure 7.**
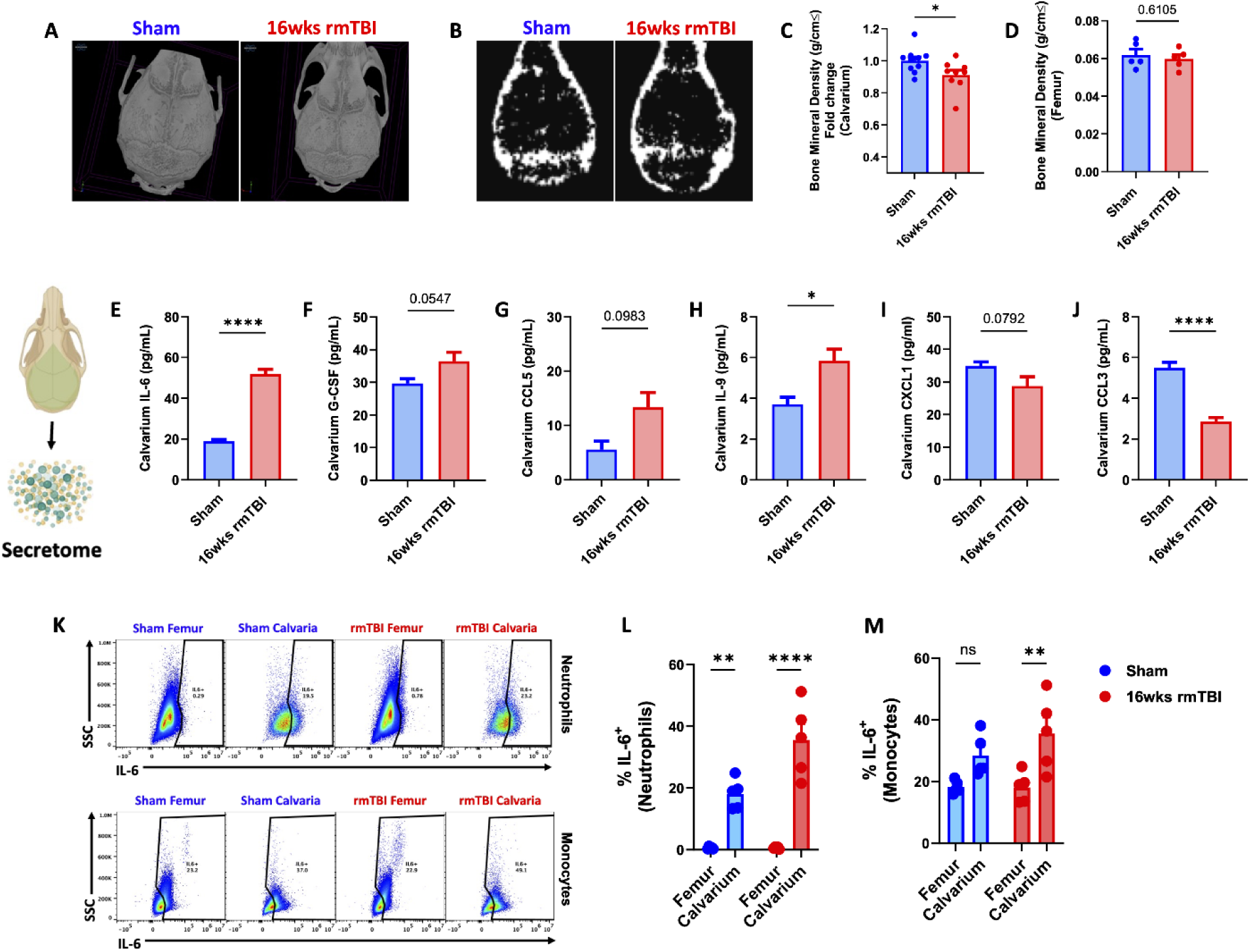
Chronic effects of rmTBI are associated with increased IL-6 secretion and decreased bone mineral density in the skull. Representative microCT images of skull calvarium in sham and rmTBI mice after 16-weeks show no overt fracturing (**A**). Representative DEXA scan images (**B**) depicting bone mineral density after rmTBI were quantified (**C**). Cytokine assessment of the skull calvaria secretome was examined by multiplex ELISA and the most significant alterations are shown (**E-J**). rmTBI-induced IL-6 secretion was confirmed by flow cytometric evaluation of IL-6 production in skull versus femur myeloid cells (**K-M**). Data were analyzed using Student’s T-test (**C-J**) or two-way ANOVA with Tukey’s post-hoc test for multiple comparisons (**L-M**). \*\*\*\**P* < 0.0001, \*\**p* < 0.01, and \**p* < 0.05. rmTBI: repeated mild traumatic brain injury, cm centimeter, g gram, pg picogram, ml milliliter, ns not significant, SSC side scatter, wk weeks.

### rmTBI reprograms the skull bone marrow secretome toward metabolic dysfunction and senescence-associated pathways

Bidirectional skull-brain communication may be mediated by secretory factors. To further elucidate the protein factors released by the skull after rmTBI, we profiled the calvarial secretome using mass spectrophotometry-based proteomics. Separation of sham and rmTBI groups by principal component analysis (PCA) revealed a distinct global shift in the proteomic secretome of bone marrow cells following trauma (**Figure 8A**), consistent with widespread functional reprogramming. Volcano plot of up- and down-regulated proteins showed significant increases in Sag, Atg12, Sh3bp1, Cplx3, Slc4a10, and Commd1b protein abundance after rmTBI, including dramatic decreases in S100a5 and Omp (**Figure 8B**). Heat map analysis of differentially regulated proteins (**Figure 8C**) revealed significant increases in proteins associated with apoptosis (Anxa5), ECM remodeling/fibrotic responses (Mcpt4, Ace), stress-induced transcriptional reprogramming (Hmga1, Brd4), and lipid metabolism (Abca8b). Proteins down-regulated after rmTBI include those associated with cytoskeleton and structural regulation (Dmd, Was, Tbca) and energy and metabolic pathways (Gapdh, Acat1, Vdac1, Casq1). Reactome pathway analysis revealed significant decreases in the citric acid TCA cycle, respiratory electron transport, glucose and pyruvate metabolism, glycolysis, and mitochondrial biogenesis after 16-weeks of rmTBI (**Figure 8D**). Reactome pathways that were increased after rmTBI included DNA double strand break response, DNA damage telomere stress-induced senescence, and DNA double strand break repair. Heat map analysis of Reactome pathways related to mitochondrial metabolism (Respiratory electron transport, Complex I biogenesis, Cristae formation, Gluconeogenesis, and Regulation of pyruvate metabolism) show widespread down-regulation of proteins involved in metabolic function (e.g., Nduf family, Atp5 subunits, Gapdh) (**Figures 8E-I**). Taken together, our findings suggest that the accumulative effects of rmTBI induce a skull secretome associated with metabolic dysfunction and senescence.

**Figure 8.**
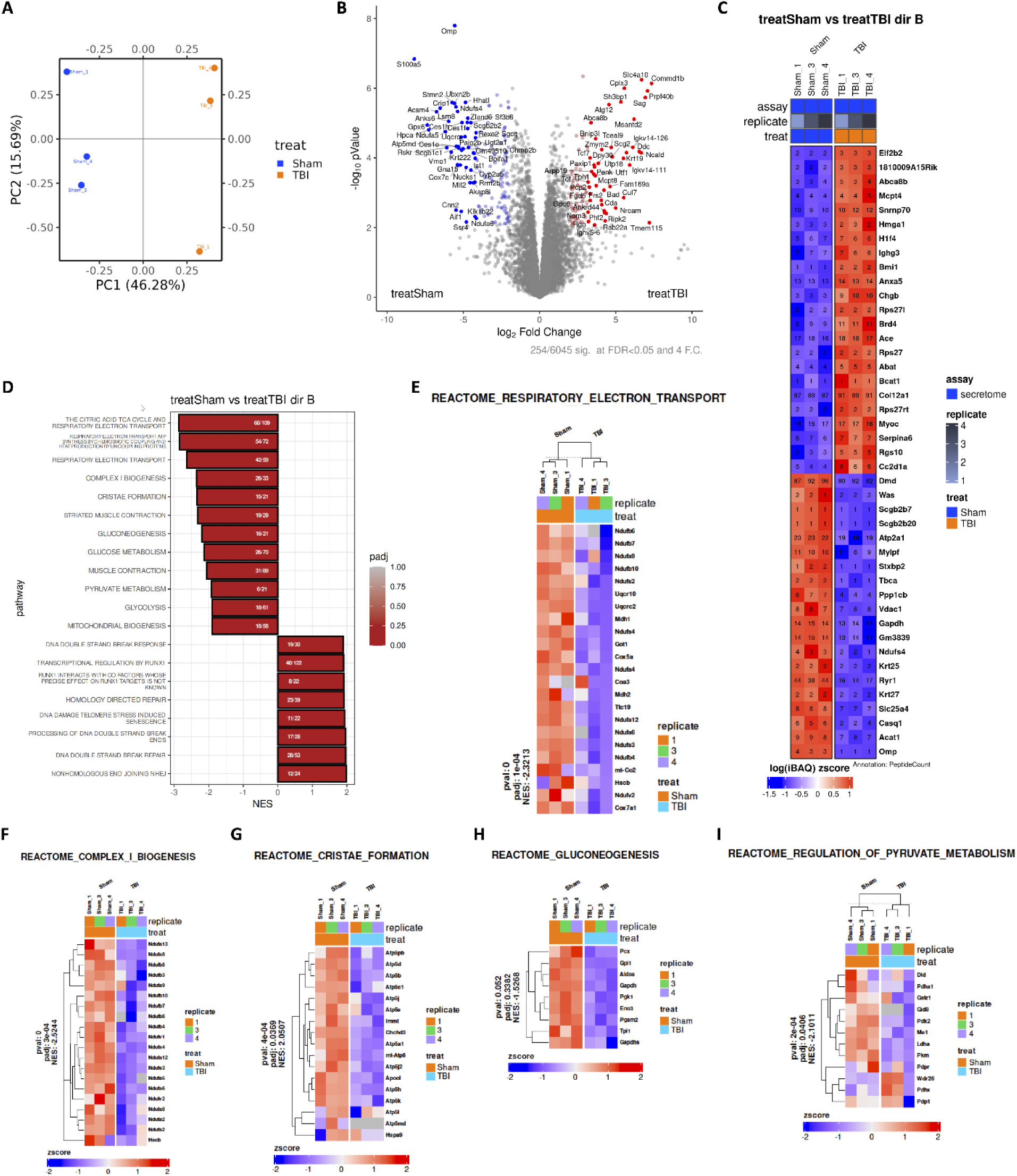
Proteomic analysis of the skull secretome after rmTBI reveals metabolic dysfunction and senescence-related pathways. Principal component analysis (PCA) of mass spectrometry-based proteomic profiling demonstrates clear separation between sham and rmTBI calvarial bone marrow secretomes (N=3/grp), indicating global remodeling of secreted protein composition following repeated injury (**A**). Volcano plot highlighting significantly up- and down-regulated proteins in the rmTBI skull secretome relative to sham, including increased abundance of Sag, Atg12, Sh3bp1, Cplx3, Slc4a10, and Commd1b, and marked reductions in S100a5 and Omp (**B**). Heat map of differentially expressed proteins reveals enrichment of factors associated with apoptosis (Anxa5), extracellular matrix remodeling and fibrotic responses (Mcpt4, Ace), stress-induced transcriptional regulation (Hmga1, Brd4), and lipid metabolism (Abca8b), alongside suppression of proteins involved in cytoskeletal organization (Dmd, Was, Tbca) and energy metabolism (Gapdh, Acat1, Vdac1, Casq1) (**C**). Reactome pathway analysis identifies significant down-regulation of pathways related to mitochondrial metabolism, including the tricarboxylic acid (TCA) cycle, respiratory electron transport, glycolysis, glucose and pyruvate metabolism, and mitochondrial biogenesis following 16 weeks of rmTBI (**D**). In contrast, pathways related to DNA double-strand break response, telomere stress-induced senescence, and DNA repair are significantly enriched. Heat map visualization of Reactome pathways associated with mitochondrial function-including respiratory electron transport (**E**), Complex I biogenesis (**F**), cristae formation (**G**), gluconeogenesis (**H**), and regulation of pyruvate metabolism (**I**)-demonstrates coordinated suppression of metabolic proteins such as NDUF family members, ATP5 subunits, and GAPDH. NES, normalized enrichment score, pval p value, padj adjusted p value.

### rmTBI skull secretome induces neurometabolic stress and reduces energetic reserve in cortical and hippocampal tissue

To determine whether factors released from rmTBI-exposed skull bone marrow directly impact brain energy metabolism, cortical and hippocampal tissue punches were stimulated with artificial cerebrospinal fluid (aCSF), sham skull secretome, or rmTBI skull secretome and analyzed using Seahorse extracellular flux assays (**Figure 9A**). In the hippocampus, exposure to the TBI skull secretome significantly increased basal oxidative phosphorylation compared to aCSF controls, while maximal oxidative capacity remained unchanged, indicating preserved mitochondrial respiratory capacity (**Figures 9B-C**). Despite this, spare respiratory capacity was significantly reduced in both cortical and hippocampal tissue exposed to the TBI secretome relative to aCSF, reflecting diminished metabolic flexibility (**Figure 9D**).

**Figure 9.**
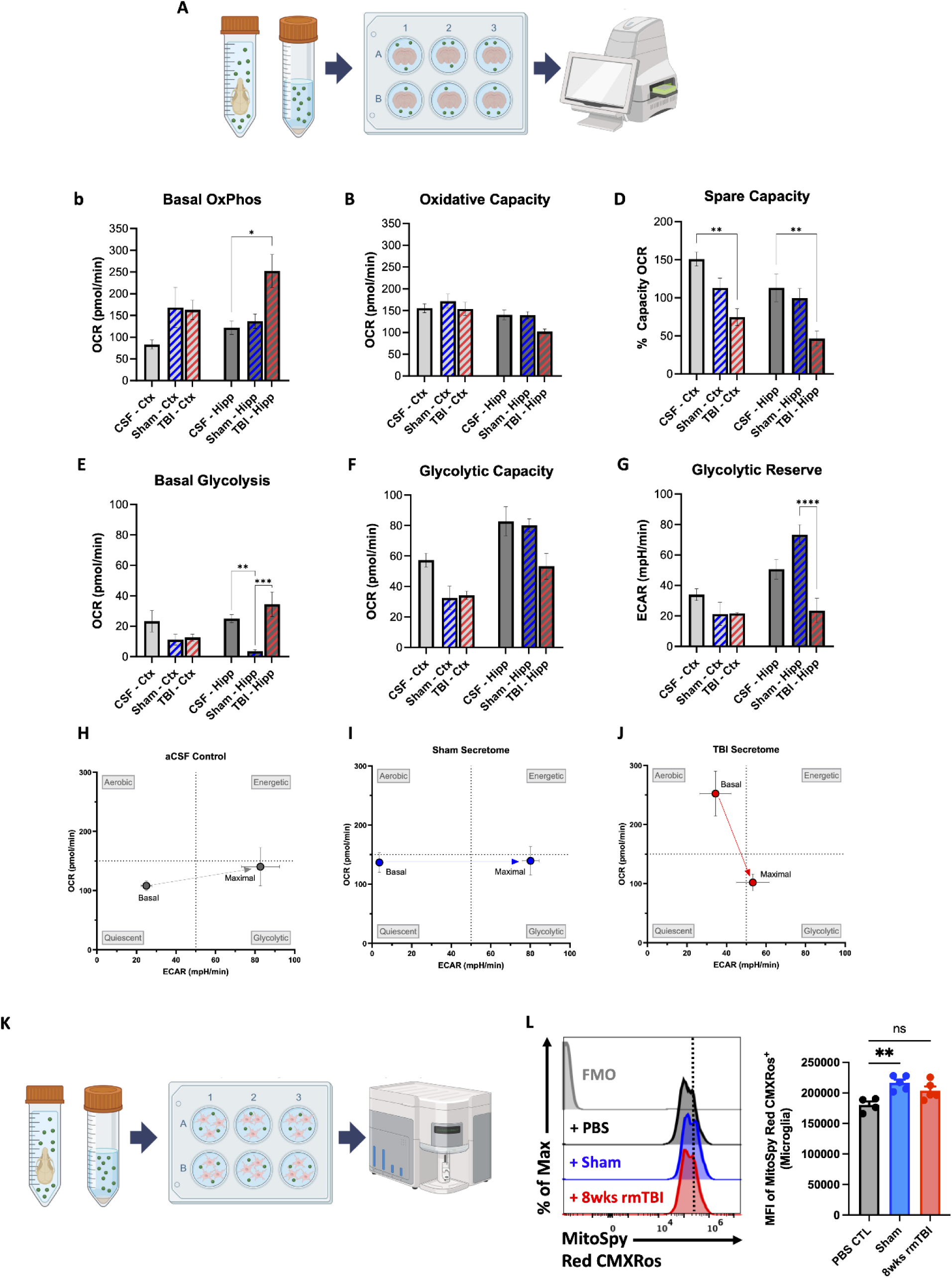
rmTBI skull secretome induces neurometabolic stress and reduces energetic reserve in brain tissue and microglia. Experimental schematic depicting ex vivo stimulation of cortical and hippocampal tissue punches with artificial cerebrospinal fluid (aCSF), sham skull secretome, or rmTBI skull secretome followed by Seahorse extracellular flux analysis (**A**). Seahorse analysis of oxidative metabolism demonstrates that rmTBI skull secretome significantly increases basal oxidative phosphorylation in hippocampal tissue relative to aCSF controls (**B**), while maximal respiratory capacity remains unchanged (**C**). Spare respiratory capacity is significantly reduced in both cortex and hippocampus following rmTBI secretome exposure, reflecting diminished metabolic flexibility (**D**). Glycolytic analysis reveals region-specific effects, with sham hippocampal tissue exhibiting reduced basal glycolysis relative to aCSF controls and rmTBI secretome inducing a significant increase in basal glycolysis compared to sham (**E**). Glycolytic capacity shows a non-significant downward trend in both cortex and hippocampus following rmTBI secretome stimulation (**F**). Glycolytic reserve is significantly reduced in hippocampal tissue exposed to rmTBI skull secretome compared with sham (**G**). Energy phenotype mapping (OCR vs ECAR) reveals that aCSF-(**H**) and sham-stimulated (**I**) tissue shifts toward a more energetic/glycolytic phenotype, whereas rmTBI skull secretome induces a downward and leftward shift toward a quiescent metabolic state (**J**), consistent with suppressed bioenergetic engagement. Experimental schematic depicting ex vivo stimulation of freshly-isolated microglia with PBS (control), sham skull secretome, or rmTBI skull secretome followed by flow cytometry analysis (**K**). Sham skull secretome significantly increases mean fluorescence intensity of MitoSpy Red CMXRos relative to PBS controls, an effect that is absent following rmTBI skull secretome stimulation (**L**). Data were analyzed using t-test and one-way ANOVA with Tukey’s post-hoc test for multiple comparisons. \*\*\*\**p* < 0.0001, \*\*\**p* < 0.001, \*\**p* < 0.01, and \**p* < 0.05. rmTBI: repeated mild traumatic brain injury, Ctx cortex, Hipp hippocampus, ECAR extracellular acidification rate, OCR oxygen consumption rate, mpH/min milli-pH units per minute, max maximum, MFI mean fluorescence intensity, SSC side scatter, wk weeks.

Analysis of glycolytic parameters revealed region-specific effects. Sham hippocampal tissue exhibited significantly lower basal glycolysis compared to aCSF controls, whereas TBI hippocampal tissue displayed a significant increase in basal glycolysis relative to sham (**Figure 9E**). In contrast, glycolytic capacity showed a downward trend in both cortex and hippocampus following TBI secretome exposure, although this did not reach statistical significance (**Figure 9F**). Notably, glycolytic reserve was significantly reduced in the hippocampus exposed to th e TBI skull secretome compared to sham (**Figure 9G**).

Energy phenotype analysis (OCR vs ECAR) was then used to assess shifts in basal metabolic state. aCSF and sham secretome-stimulated brain tissue revealed a shift toward a more energetic/glycolytic phenotype (**Figures 9H-I**). In contrast, rmTBI secretome-stimulated brain tissue showed a shift downward and leftward, moving away from the energetic/aerobic quadrants toward a more quiescent state (**Figure 9J**). Seahorse energy phenotype mapping demonstrated a transmissible shift toward a quiescent metabolic phenotype, consistent with suppressed bioenergetic activity and impaired metabolic engagement after rmTBI.

Finally, to assess whether the rmTBI skull-derived secretome altered mitochondrial membrane potential in otherwise healthy young adult microglia (**Figure 9K**). Interestingly, sham skull secretome induced a significant elevation in mitochondrial membrane potential relative to PBS control (p<0.01), whereas this effect was suppressed in the rmTBI skull secretome-stimulated group (**Figure 9L**).

Collectively, these findings demonstrate that the rmTBI skull secretome induces a state of neurometabolic stress characterized by elevated baseline energy utilization, preserved maximal respiratory capacity, and marked reductions in both oxidative and glycolytic reserve in cortical and hippocampal tissue. This metabolic inflexibility is further reflected at the cellular level, as rmTBI skull-derived factors suppress mitochondrial membrane potential in microglia, in contrast to the stimulatory effects observed with sham secretome. Together, these data indicate that rmTBI-conditioned skull secretomes transmit a bioenergetically constraining signal to the brain, promoting a shift toward a quiescent metabolic state and rendering both neural tissue and resident immune cells more vulnerable to additional energetic stress. These findings identify skull-derived factors as potent modulators of brain bioenergetic homeostasis following rmTBI.

### Regional microglial activation phenotypes after 16-weeks of rmTBI

To confirm that our 16-week rmTBI paradigm with inter-session recovery induces chronic microglial activation phenotypes, we performed flow cytometry using sham and rmTBI brains. To increase spatial understanding, we analyzed changes in cortex, hippocampus, and striatal regions. A global increase in microglial proliferation, as measured by hemispheric cell count, was determined using two-way ANOVA group analysis (p=0.01, **Figure 10A**). Expression of the disease-associated microglia (DAM) marker, Clec7a, was significantly upregulated in the cortex after rmTBI (**Figure 10B**). Production of the pro-inflammatory cytokine, TNF, was regionally elevated in the cortex and hippocampus, but not striatum (p=0.008, **Figure 10C**). Interestingly, we found an increase in myelin CNPase antigen levels inside microglia in the cortex (p<0.05) and striatum after rmTBI, and decreased intracellular detection in hippocampal microglia, resulting in a statistical interaction between rmTBI and brain region by two-way ANOVA group analysis (p<0.0001, **Figure 10D**). Lastly, we found that SA-βGal activity was significantly increased in striatal microglia (p<0.05), but not cortical microglia (**Figure 10E**). These findings highlight the regional heterogeneity of microglial responses to rmTBI, with increased Clec7a expression and myelin engulfment seen predominantly in cortical microglia, increased SA-βGal activity largely restricted to striatal microglia, and injury-induced proliferation seen across all regions.

**Figure 10.**
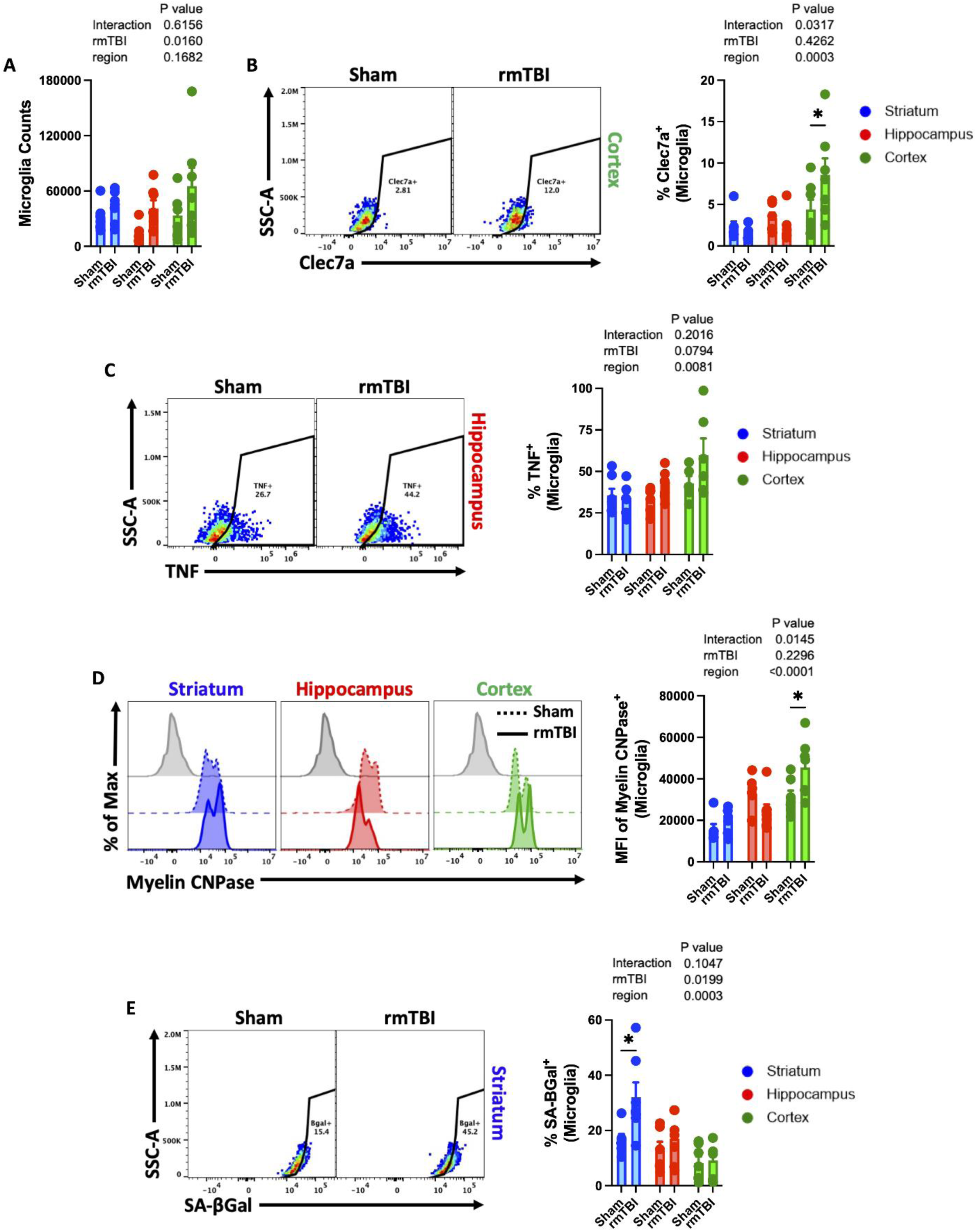
Regional microglial activation phenotypes after 16-weeks of rmTBI. Assessment of microglia activation was performed on cortical, hippocampal, and striatal tissue via flow cytometry. Counting bead-estimated live CD45^int^CD11b^+^Ly6C^-^ microglial counts were quantified after 16-weeks of rmTBI (**A**). The percentage of Clec7a-positive microglia is shown across regions (**B**). TNF production in microglia was quantified as shown (**C**). Representative histograms reveal the relative level of intracellular myelin CNPase antigens detected in microglia after rmTBI (**D**). Representative dot plots show the frequency of microglia with senescence-associated beta-galactosidase activity after rmTBI (**E**). For all histograms, light gray = fluorescence minus one (FMO) control. Data were analyzed using two-way ANOVA group analysis with Tukey’s post-hoc test for multiple comparisons (**A-E**). \**P* < 0.05. rmTBI: repeated mild traumatic brain injury, max maximum, SSC-A side scatter-area, wk weeks.

## Discussion

This study identifies a previously unrecognized pathophysiological cascade linking rmTBI to chronic systemic inflammation, innate immune dysfunction, and features of accelerated aging. While prior work has shown infiltration of skull bone marrow-derived leukocytes into the brain after moderate-to-severe TBI[43], our findings indicate that in mild injury paradigms-where overt leukocyte infiltration is absent-long-term neurological consequences are driven instead by hematopoietic stem/progenitor cell dysfunction, impaired leukopoiesis, and altered skull–brain communication. We demonstrate that each head impact induces a transient wave of LSK cell proliferation and de novo monocyte production within the bone marrow; however, repeated proliferative cycling over weeks of rmTBI leads to telomere shortening, early replicative senescence, and permanent cell-cycle arrest. This culminates in reduced leukocyte output and leukopenia in otherwise young adult mice. Moreover, rmTBI reprograms the inflammatory landscape of the skull bone marrow, characterized by elevated IL-6 and a senescence-associated proteomic secretome enriched for pathways related to metabolic dysfunction and DNA damage responses. Importantly, we show that skull-derived secretory factors are sufficient to disrupt mitochondrial metabolism in brain tissue and microglia, providing functional evidence for skull-brain signaling as a mediator of chronic neuroimmune and neurometabolic dysfunction following repeated mild head injury.

Recent groundbreaking work by Kolabas et al., demonstrated molecular heterogeneity in bone marrow compartments across the body[44]. The skull has a unique molecular and functional composition that can react and respond to underlying brain pathologies. In our study, we evaluated bone marrow responses to rmTBI in skull and femur. The former is a distally-located long, thick bone rich in marrow, while the latter is a thin flat bone with a spongy diploë center containing sparse marrow. Despite these fundamental differences, the hematopoietic response in both bones were similarly sensitive to the initial rmTBI. Surprisingly, unlike femur, skull bone marrow cells appeared refractory to the second week of rmTBI. The reason for this blunted response is unclear, but could include innate immune training, a tolerance mechanism, immune suppressive signals, or metabolic disruption from the physical force of the impact. Our findings did show that mitochondrial membrane potential of skull LSKs decreased significantly after rmTBI, suggesting ATP production and overall cellular activity could be diminished. Skull bone marrow cells residing in their diploë microenvironment may likewise be subject to a metabolic crisis caused by the immediate physical forces applied to the bone during a TBI. Due to cell-intrinsic mechanisms caused by direct ‘trauma’, and/or cell-extrinsic mechanisms mediated by trauma-induced factors, a senescence phenotype was induced. Indeed, each head impact caused a wave of proliferation in femur LSKs, ultimately leading to replicative senescence by 16 weeks of rmTBI. However, skull LSKs did not show the same recurrent response. Nevertheless, after 16 weeks of rmTBI skull LSK cells displayed a senescent phenotype characterized by reduced proliferative capacity and higher SA-βGal activity. Whether this was attributed to direct injury, chronic stress, bystander senescence, or replicative senescence is not known. However, we cannot rule out the latter, because the proliferative status of skull LSKs beyond 24 hrs post-rmTBI at week 16 was not measured. Indeed, telomere lengths were significantly reduced in both bone marrow compartments, with femur showing a near ∼50% reduction after rmTBI compared to sham control. These findings are consistent with previous studies indicating shorter blood leukocyte telomere lengths were associated with higher total symptom burden at 3-months following mild TBI[45], and may be accelerated in older trauma patients[46]. Shorter telomere lengths were also found in saliva of collision sports athletes independent of concussion history or sex[47]. Telomere length is a predictive biomarker for injury prognosis in juvenile rats following a concussion[48], and shorter telomeres have been linked to higher risk of age-related brain diseases in humans[49, 50]. Whether this association is causal remains to be seen, but our prior work suggests that TBI initiates bone marrow dysfunction which can, in turn, independently drive neurological dysfunction[22].

Given its close proximity to the brain, the skull bone marrow compartment, and the soluble factors it releases, may be key mediators of secretory communication. The recent discovery of synaptic proteins in the human skull suggests that communication along the skull-meninges-brain axis may be bidirectional[44, 51]. However, cytokine and proteomic profiling of the skull secretome in the context of TBI has not been reported. It stands to reason that repeated head impacts directly affect immune signaling pathways in bone marrow cells which contribute to an altered inflammatory milieu. Senescent cells often adopt a senescence-associated secretory phenotype (SASP), consisting of a pro-inflammatory, pro-remodeling mixture of factors (e.g., cytokines, chemokines, growth factors, proteases, and extracellular matrix-modifying proteins)[52, 53]. IL-6 is one of the core SASP factors most commonly observed across cell types and models[54, 55]. Among the 32 cytokines measured in the multiplex ELISA panel, IL-6 exhibited the greatest increase in secretion after 16-weeks of rmTBI. IL-6 is one of the earliest and most robustly upregulated cytokines in the injured brain[56, 57]. Its effects are context-dependent, acting as both a pro-inflammatory and neurotrophic factor depending on timing, location, and cell type involved[58]. Interestingly, rmTBI elicited higher IL-6 production in skull myeloid cells compared to femur, suggesting the skull compartment may represent a regionally specialized immune niche that contributes disproportionately to rmTBI-associated neuroinflammation. IL-6 can also decrease bone mineral density by promoting bone resorption and impairing bone formation, especially in chronic inflammatory states[59, 60]. This may partially explain the decreased bone mineral density seen in the skull at 16 -weeks post-rmTBI. Moreover, this may also explain why righting reflex times show a delayed increase compared to sham control at later timepoints. Whether this cytokine diffuses into the brain and what role, if any, it may have on neurological outcome remains to be investigated.

Our proteomic survey of the skull (i.e., calvarial) secretome yielded several surprising and novel findings. rmTBI skull-derived secretome proteomics revealed coordinated metabolic and stress-response remodeling. Hallmark analysis showed decreased KRAS signaling and increased DNA repair, while GO and Reactome pathways indicated broad suppression of mitochondrial respiration (respiratory chain, NADH dehydrogenase, cytochrome, Complex I biogenesis, cristae formation) and core metabolic flux (gluconeogenesis, pyruvate metabolism). In contrast, vesicle-coat proteins were elevated, suggesting enhanced extracellular vesicle trafficking. Several DNA repair pathways were upregulated, although processing of double-strand breaks appeared attenuated. This profile is consistent with a cellular stress or senescence-like state, where cells suppress energy-intensive anabolic programs and activate repair and maintenance mechanisms. Collectively, these findings indicate a shift toward a hypometabolic, stress-adapted, and vesicle-secretory phenotype, consistent with cellular senescence or chronic injury adaptation. The functional significance of these secreted factors was confirmed. At the cellular level, rmTBI skull-derived factors suppressed mitochondrial membrane potential in microglia. At the tissue level and across regions, these factors induced a bioenergetic profile that closely resembles senescence-associated metabolic inflexibility, in which cells remain metabolically active but lack the capacity to respond to additional energetic demands. Together with evidence of telomere shortening, SA-βGal induction, and a pro-inflammatory secretome, these findings suggest that rmTBI promotes a senescence-like metabolic state in brain tissue rather than overt mitochondrial failure.

Microglia play diverse roles in maintaining brain health and responding to injury. Regional microglial heterogeneity has been observed in animal models of rmTBI and in humans[61–63]. Indeed, different brain regions exhibit distinct microglial activation profiles and transcriptional signatures following mild injury. However, region-specific functional responses to TBI are less well understood. Our study revealed some surprising findings, including evidence for an increased senescent-like phenotype (i.e., SA-βGal activity) in striatal microglia after rmTBI. Several rodent models of mTBI show microglial reactivity in the striatum, especially when examining deeper subcortical structures over time. The striatum has dense synaptic innervation, which require microglia to be tightly coupled to synaptic maintenance and pruning. The striatum also has a unique vascular architecture and high metabolic demand, which may be disturbed after TBI, affecting glial metabolic coupling. Nevertheless, it is not clear why microglia in deeper brain regions would be more prone to premature senescence. Our finding that cortical microglia upregulate the DAM marker, Clec7a, is expected given their proximity to the impact site (i.e., calvarium). We have previously reported that microglia adopt a pro-phagocytic phenotype late after a single moderate-to-severe TBI, characterized by increased engulfment of myelin and apoptotic neurons[64]. We found a similar phenotype in cortical microglia after rmTBI. However, it is unclear whether the increase in myelin removal is required for debris clearance and injury resolution, or pathological in nature. A deeper understanding of this functional heterogeneity will be critical for targeting region-specific populations of neurotoxic microglia.

Given the profound immunological repercussions of rmTBI seen in our hands, it is reasonable to assume our 16-week repeated weight drop model causes moderate-to-severe injury. However, the injury parameters were carefully titrated to induce a mild brain injury sufficient to trigger subtle pathophysiological responses without causing extensive damage. As such, no overt subdural bleeding, skull fracturing, brain infiltration of leukocytes, body weight loss, or mortality was observed. No difference in righting reflex time was noted until after 11-weeks post-rmTBI, suggesting the accumulated stress of prior head impacts reached a pathological threshold at this time. Moreover, we would argue that the severity of behavioral impairment at 16-weeks post-rmTBI was also relatively mild across tests. Thus, it appears that even low-intensity impacts in younger animals can have a compounding effect on bone marrow aging and peripheral immunity.

This study is not without limitations. Due to concerns for animal welfare, female mice, which are comparably smaller and potentially more vulnerable to our rmTBI model were not included in our experimental design. However, several studies indicate younger female athletes may have more enduring negative symptoms associated with sports-related mild concussive injury[65, 66]. We also acknowledge that the longitudinal timeline of our study (e.g., 1 −2-week, 8-week, and 16-week endpoints) does not adequately capture the sequential timing of events. For example, it is not entirely clear whether the onset of bone marrow senescence induction precedes neuroinflammation or neurological impairment. As such, a causal association cannot be established. Finally, we recognize this study is largely descriptive. Given the potential implications of the findings, the study was published ahead of full mechanistic resolution, which remains a focus of ongoing work.

In conclusion, our findings suggest mild sub/concussive injury directly impacts skull bone marrow stem/progenitor cells, hematopoiesis, and peripheral immunity. Normal age-related replicative senescence was accelerated with each subsequent head impact, even in distal bone marrow compartments. Early induction of hematopoietic stem/progenitor cell senescence results in leukopenia, innate immune dysfunction, and an inflammatory cranial milieu with microglia-activating potential. Importantly, factors secreted from the skull after rmTBI could induce metabolic dysfunction, providing a neural basis of neurological impairment in mice with otherwise mild neuropathology. Although diminished immune-mediated tissue repair and regeneration can promote chronic inflammation, lead to loss of CNS homeostasis, neurological impairment, and increased risk/severity of infection, our key finding that TBI is a chronic immunodegenerative disease has yet to be fully validated in humans. This study warrants further investigation into the immune status of individuals with a history of repeated mild head impacts/concussions.

## Acknowledgments

PD conducted all the behavioral experiments, carried out the statistical analyses, and wrote the manuscript. RK, TD, BW, SM, GG, JK, KG, and CT assisted with the experiments, figures, and samples. PD performed the weight drop TBI model. JA and SM assisted with the microCT imaging. AM conducted the MS-based proteomic analysis. EU performed the Seahorse metabolism experiments. KB, RF, MM, and SM for their assistance in developing and obtaining veterinary approval for the injury model used in this study. RR conceived and designed the project. RR and RK edited the manuscript. Schematics were made using BioRender. We thank Katherine Brannick, DVM, for her expert veterinary oversight and invaluable assistance with the monitoring and assessment of mice used in this study. Her contributions were essential to ensuring animal welfare.

---

**Supplemental Figure 1.**
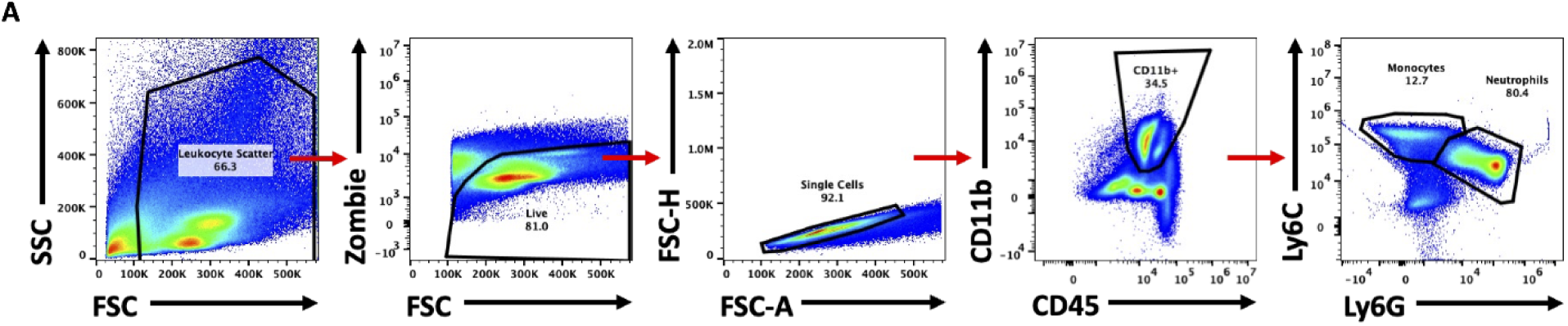
Bone marrow cell gating strategy. Viable singlet bone marrow leukocytes were identified using scatter properties and Zombie dye exclusion, followed by gating on CD45⁺CD11b⁺ myeloid cells. Monocytes were defined as Ly6C^+^ Ly6G⁻ cells, and neutrophils as Ly6G⁺ cells. SSC side scatter, FSC-A forward scatter-area.

